# Structure of the endosomal CORVET tethering complex

**DOI:** 10.1101/2023.10.13.562242

**Authors:** Dmitry Shvarev, Caroline König, Nicole Susan, Lars Langemeyer, Stefan Walter, Angela Perz, Florian Fröhlich, Christian Ungermann, Arne Moeller

## Abstract

Cells depend on their endolysosomal system for nutrient uptake and downregulation of plasma membrane proteins. These processes rely on endosomal maturation, which requires multiple membrane fusion steps. Early endosome fusion is promoted by the Rab5 GTPase and its effector, the hexameric CORVET tethering complex, which is homologous to the lysosomal HOPS. How these related complexes recognize their specific target membranes remains entirely elusive. Here, we solved the structure of CORVET by cryo-electron microscopy and revealed its minimal requirements for membrane tethering. As expected, the core of CORVET and HOPS resembles each other. However, the function-defining subunits show marked structural differences. Notably, we discovered that unlike HOPS, CORVET depends not only on Rab5 but also on phosphatidylinositol-3-phosphate (PI3P) and membrane packaging defects for tethering, implying that an organelle-specific membrane code enables fusion. Our data suggest that both shape and membrane interactions of CORVET and HOPS are conserved in metazoans, thus providing a paradigm how tethering complexes function.

**One-Sentence Summary:** Using cryo-EM, we solved the structure of the CORVET tethering complex and explained how it promotes endosome fusion.

## Introduction

Eukaryotic cells maintain an elaborate endomembrane system of organelles. This interconnected network depends on the vesicular transport and relies on conserved types of machinery for vesicle generation at the donor organelle and fusion at the acceptor organelle (Borchers et al., 2021; Gomez-Navarro and Miller, 2016; Rout and Field, 2017). For proper intracellular recognition, each organelle exhibits distinct membrane compositions and shapes. Identity markers include specific phosphoinositides (PIPs) in the lipid bilayer (Posor et al., 2022) and distinct peripheral membrane proteins such as Rab GTPases (Rabs) (Borchers et al., 2021; Goody et al., 2017). Together, these elements guide trafficking effector proteins to their specific membrane and enable the effective direction of vesicles to their acceptor organelles (Bigay and Antonny, 2012; Herrmann et al., 2023; Posor et al., 2022; Wilmes and Kümmel, 2023).

Rabs are an integral part of the membrane fusion cascade, as their depletion leads to impaired membrane trafficking (Singer-Krüger et al. 1994, Wichmann et al. 1992). As switch-like proteins, Rabs only interact with their effector proteins upon activation by guanine nucleotide exchange factors (GEFs) and binding to GTP while requiring a GTPase-activating protein (GAP) to hydrolyze GTP for their inactivation (Barr, 2013; Borchers et al., 2021; Goody et al., 2017; Hutagalung and Novick, 2011; Wandinger-Ness and Zerial, 2014).

The most prominent effector proteins are tethering and fusion factors and complexes (Beek et al., 2019; Spang, 2016; Ungermann and Kümmel, 2019; Wickner and Rizo, 2017; Zhang and Hughson, 2021). They tether the membranes and recruit Sec1/Munc18 proteins (SM proteins) to promote the zippering of SNAREs from each membrane into a four-helix bundle to trigger fusion (Risselada and Mayer, 2020; Ungermann and Kümmel, 2019; Wickner and Rizo, 2017; Zhang and Hughson, 2021).

The hexameric HOPS and CORVET are evolutionarily conserved tethering complexes within the endolysosomal system of eukaryotic cells (Beek et al., 2019; Kant et al., 2015; Peplowska et al., 2007; Perini et al., 2014; Seals et al., 2000; Wurmser et al., 2000). Both share four subunits. Vps11 and Vps18 form the central core to which Vps33 and Vps16 are attached as an SM module to promote SNARE assembly (Baker et al., 2015; Baker and Hughson, 2016). The two unique subunits determine the respective Rab specificity of the tethering complexes (Kant et al., 2015; Markgraf et al., 2009; Ostrowicz et al., 2010; Peplowska et al., 2007; Perini et al., 2014; Plemel et al., 2011). In CORVET, Vps3 and Vps8 bind to Rab5 on early endosomes (EE) (Markgraf et al., 2009; Peplowska et al., 2007; Perini et al., 2014), while in HOPS, Vps41 and Vps39 interact with the Rab7-like Ypt7 in yeast, and with Rab2 and lysosomal Arl8 in metazoan HOPS (Beek et al., 2019; Jiang et al., 2014; Pols et al., 2013; Seals et al., 2000; Takáts et al., 2014; Wickner and Rizo, 2017; Wilkin et al., 2008; Wurmser et al., 2000). The specificity of the distinctive subunits is fundamental to the different roles of the two complexes within the endolysosomal system. CORVET detects Rab5, which decorates early endosomes and, therefore, functions in the fusion of endocytic vesicles and among EEs. HOPS promotes the fusion of late endosomes, autophagosomes, and vacuoles. Given their central position in the endolysosomal pathway, it is not surprising that CORVET (Bayram et al., 2016) and HOPS have been linked to diseases and infection (Anderson et al., 2022; Beek et al., 2019; Miao et al., 2021; Sindhwani et al., 2017; Sofou et al., 2021; Welle et al., 2021).

So far, most mechanistic insights into endolysosomal tethers have been obtained on the yeast HOPS complex. Proteoliposome assays and *in vivo* analyses revealed how it catalyzes fusion through tethering of Ypt7-decorated membranes and promoting the assembly of membrane-anchored SNAREs (Bröcker et al., 2012; D’Agostino et al., 2017; Lürick et al., 2017; Mima et al., 2008; Stroupe et al., 2009). How CORVET works remains however unclear, and any discussions on similarity between HOPS and CORVET are speculative in the absence of structural insights.

Here, we present the structure of CORVET that resolves this puzzle. The overall structure of CORVET is conserved and highly similar to HOPS (Shvarev et al., 2022), while the Rab/membrane-interacting subunits, which control the respective functions of the two complexes, are remarkably different. Unlike HOPS, which solely depends on Ypt7 to tether membranes, CORVET requires PI3P and loose membrane packing in addition to its Rab5 GTPase for tethering. In combination, our study provides structural and functional evidence of how a multisubunit tethering complex perceives the combinatorial code of the membrane environment. As CORVET differs only at those positions from HOPS that control membrane specificity, we predict that the form and shape of the human CORVET and HOPS complexes will be similar if not identical to the yeast complexes.

## Results

### Cryo-EM structure of the CORVET complex and its specific subunits

CORVET was produced in yeast cells and isolated via a FLAG tag on Vps8 using affinity chromatography followed by size exclusion chromatography (SEC) with high purity (Figure 1A,B). The molecular weight of the purified complex measured by mass photometry (656 kDa) agreed with the predicted weight of 658 kDa (Figure 1C).

**Figure 1.**
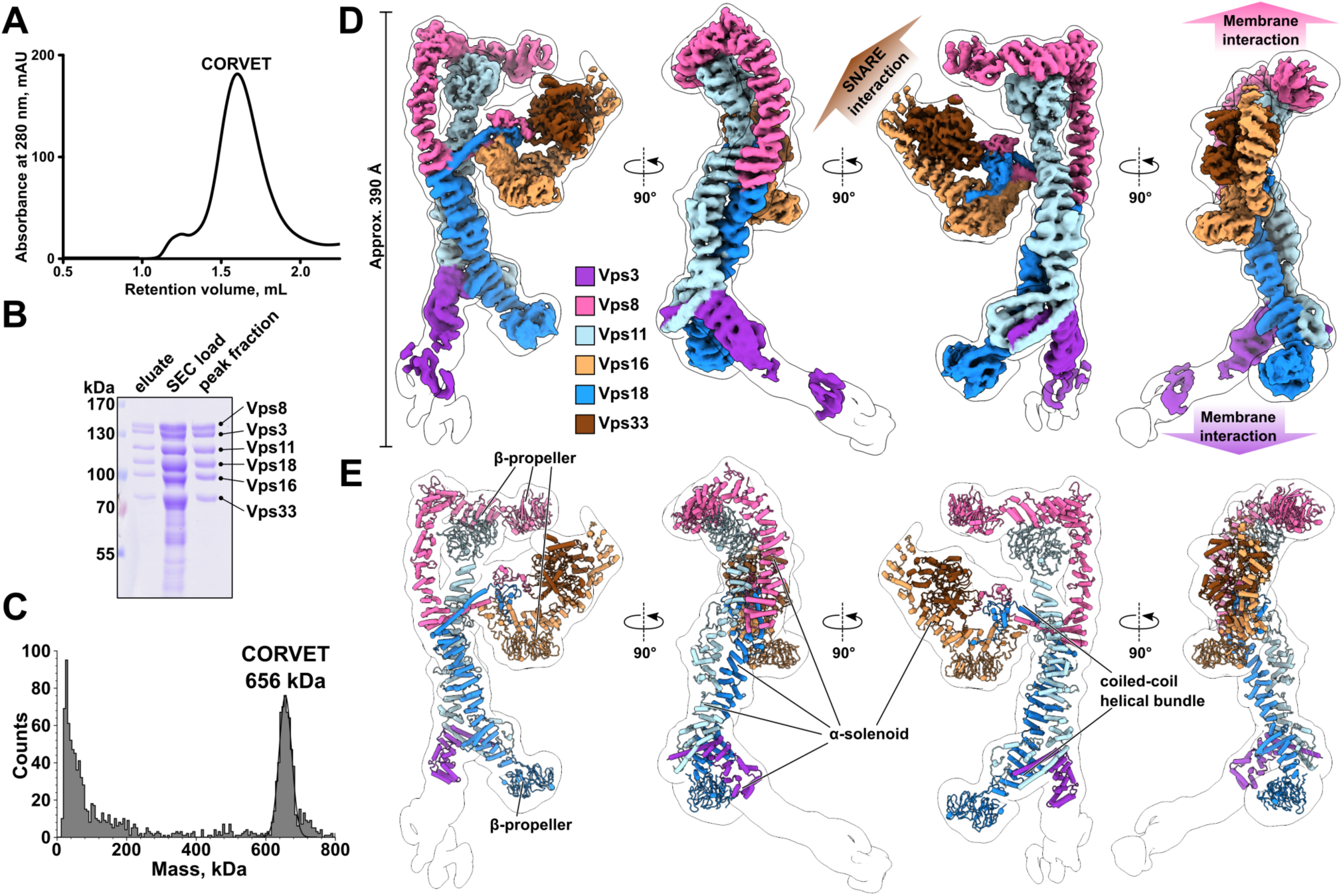
Purification and cryo-EM structure of the yeast CORVET complex. **A**, Size exclusion chromatography (SEC) of the affinity-purified CORVET. **B**, SDS-PAGE analysis of purified CORVET. Protein samples from affinity purification (eluate) and before and after SEC are shown. **C**, Mass photometry analysis of the peak fractions from SEC of CORVET. **D**, Overall architecture of the CORVET complex. Composite cryo-EM map generated from local refinement maps (see Figures S2,3) is colored by subunits assigned. The consensus maps used for local refinements is low-pass-filtered and shown as a transparent envelope with a black outline. **E,** Molecular model of CORVET fitted into the low-pass-filtered consensus map.

Negative stain electron microscopy (EM) revealed evenly distributed single particles of CORVET with no apparent aggregates in the sample (Figure S1A), allowing further single-particle cryo-EM analysis (Figures S1B, S2). From 33882 micrographs, 219391 selected particles provided a consensus map with 4.6 Å resolution. Local refinements further improved the resolution to 3.8 Å (SNARE-binding module, Figure S3, Table S3). A composite map of all local refinements (Figures 1D, S2) enabled model building of the CORVET structure (Figures 1E, 2, S4, Movie S1, Table S3).

CORVET and HOPS share strong structural homology. Their respective cores, consisting of identical subunits (Vps11 and Vps18), are virtually indistinguishable from each other (Figure 2E). A similar scenario is found for the SNARE-binding module (Vps33-Vps16) which branches off sideways from the core.

**Figure 2.**
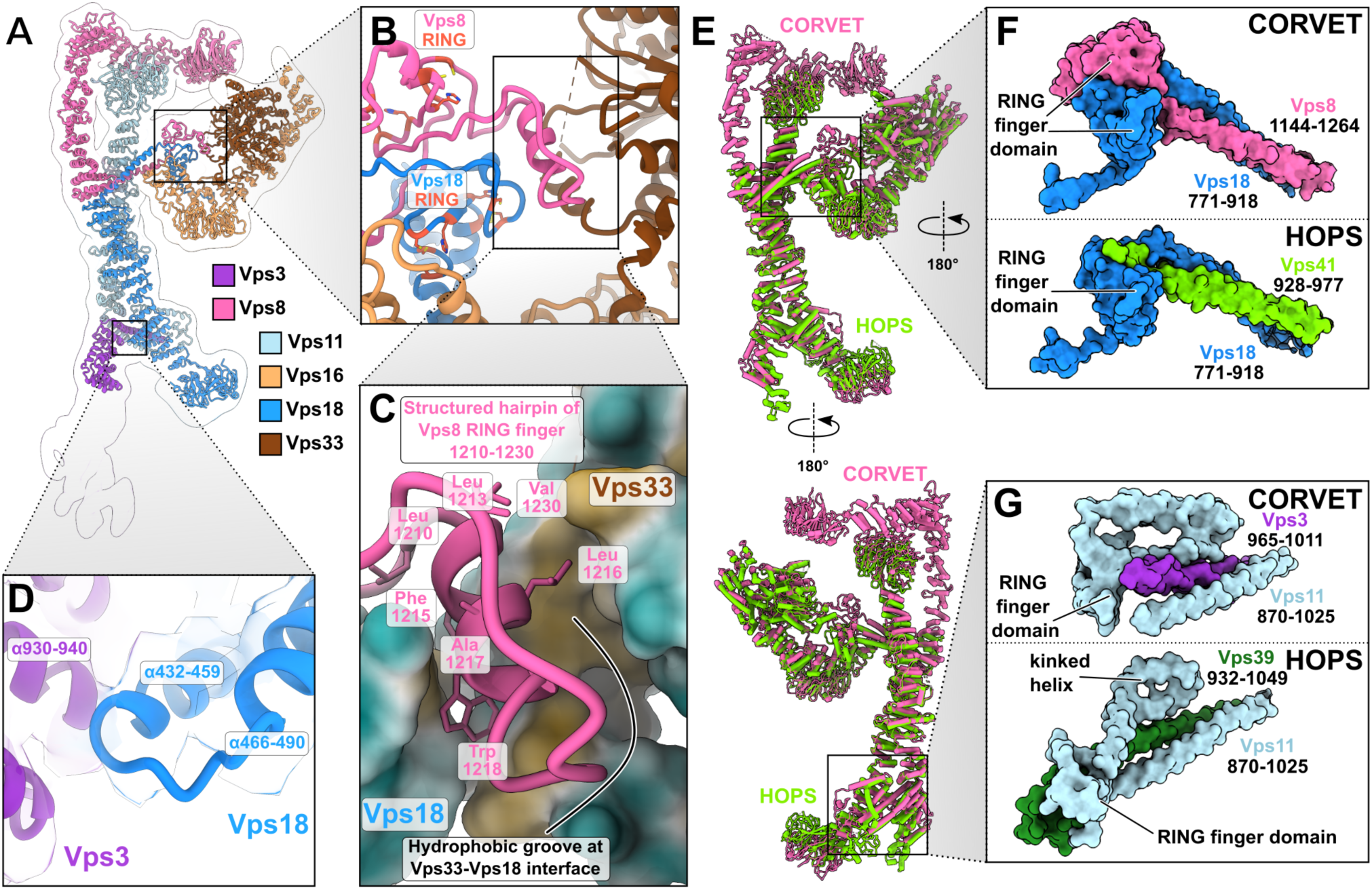
Interactions of CORVET functional subunits with the core of the complex. **A**, Model of the CORVET complex viewed from the side in ribbon representation. The low-pass-filtered consensus map is shown by a black outline. The subunits are colored as in Figure 1. **B**, Zoomed-in view of the Vps33-Vps8-Vps16 interface with Vps8 and Vps18 RING finger domains indicated. **C**, Zoomed-in view of the Vps8 structured hairpin sandwiched between Vps33 and Vps18. **D,** Interactions between the α-solenoids of Vps18 and Vps3. Associated semi-transparent cryo-EM densities are zoned around the molecular models and colored accordingly. **E,** Structural alignment of the CORVET molecular model (pink, this study) with the HOPS model (light green, PDB:7ZU0) viewed from two sides. **F,** Close-up views of C-terminal two-helix bundle and RING finger domains of CORVET Vps8 and Vps18 (top) and HOPS Vps41 and Vps18 (bottom). **G**, Close-up views of C-terminal two-helix bundle and RING finger domains of CORVET Vps3 and Vps11 (top) and HOPS Vps39 and Vps11 (bottom).

Reflecting their diverse functions within the endolysosomal system, those parts of the complexes that are responsible for target recognition show marked differences (Figure 3). Vps3 and Vps8 are specific to CORVET and located at the distal ends of the complex (Figure 1D,E). Each interlocks with the core through long C-terminal α-helices, reminiscent to the HOPS-specific Vps39 and Vps41 subunits (Figures 1E, 2E-G, 3).

**Figure 3.**
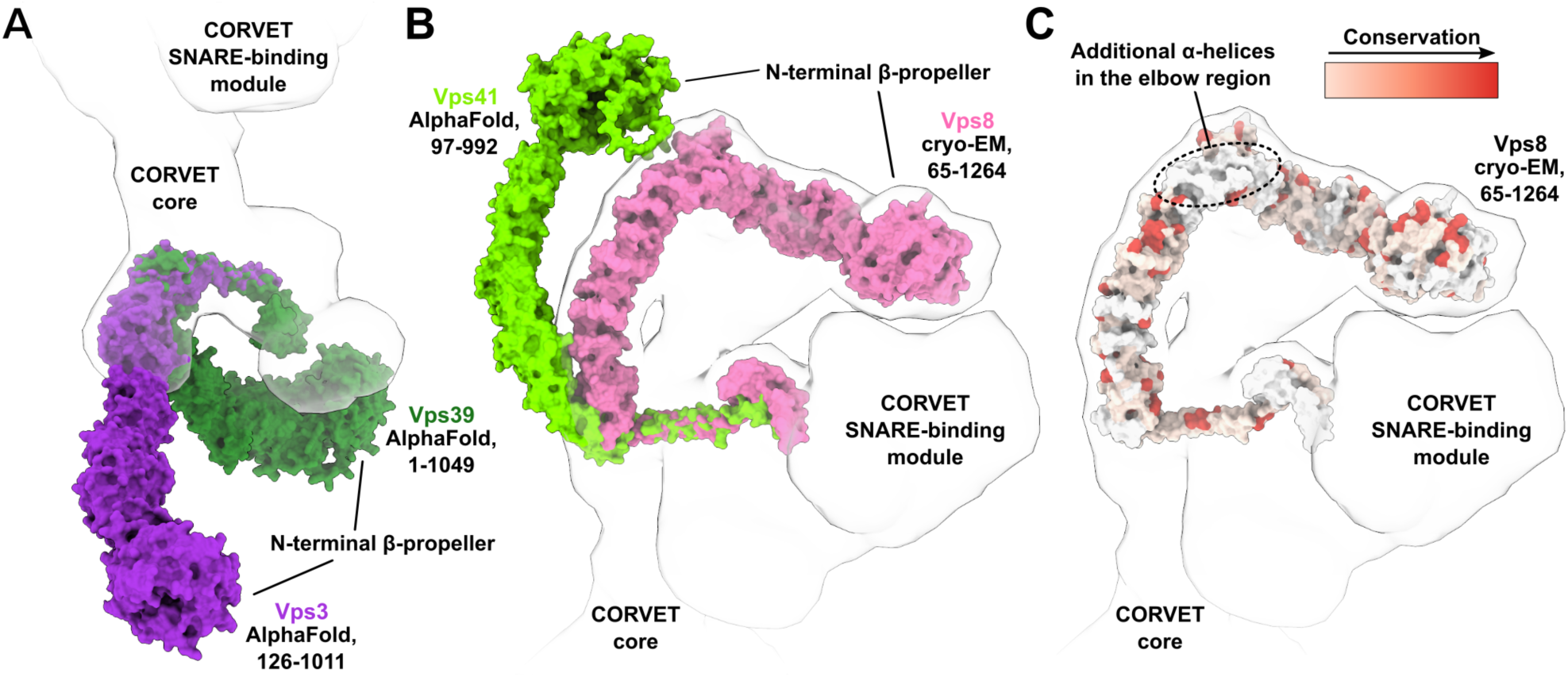
Rab-binding subunits of CORVET compared to HOPS. **A**, Structural alignment of CORVET Vps3 (violet, AlphaFold prediction) with HOPS Vps39 (dark green, AlphaFold prediction). **B,** Structural alignment of CORVET Vps8 (pink, cryo-EM structure from this study) with HOPS Vps41 (light green, AlphaFold prediction). **C**, Vps8 subunit colored by conservation according to the sequence alignment with Vps41. In **A-C**, the models of proteins are fitted into the semi-transparent volume (black outline) generated from the CORVET structure.

The extended Vps3 subunit is located at one end of the structure, but substantial flexibility in this region (Figure S5, Movies S2-4) limited the achievable resolution. As such, the N-terminal part of Vps3, carrying the β-propeller, is only visible at lower thresholds (Figure 1E, black outline). Similarly, to the analogous subunit Vps39 in HOPS, the C-terminal α-helix of Vps3 is anchored to CORVET’s core through an interaction with the RING finger domain and the long α-helix located at the C-terminus of Vps11 (Figure 2E,G). In HOPS the RING finger domain of Vps11 tightly interacts with the C-terminal portion of Vps39 that follows the long α-helix. In contrast, the CORVET subunit Vps3 lacks any further C-terminal extensions after the long α-helix, thus significantly reducing the contact area in the Vps3-Vps11 interface. Furthermore, we observe a high flexibility of Vps3 and a more distant positioning from the complex core, when compared to HOPS (Figures 1D,E, 3A). Interestingly, mutations in the Vps3-Vps11 interface can have profound impacts on the complex. As shown by mass spectrometry, CORVET variants carrying mutations in Vps11 (*vps11-1, vps11-3*) (Peterson and Emr, 2001) destabilize the complex (Figure S6A-F). In case of the *vps11-3* allele the attachment of the Vps3 subunit to the core is abolished (Figure S6C,D), whereas the *vps11-1* allele disassembles the entire complex (Figure S6E,F).

Vps8, the second Rab-binding subunit of CORVET, is located at the opposite end of the complex (Figure 1D,E). Vps8 exhibits a characteristic elbow configuration, which turns its N-terminal half towards the β-propeller of Vps11. In the equivalent Vps41 subunit of HOPS, no such coordination is observed (Figure 3B). In fact, Vps41 is curved in the opposing direction and exhibits higher flexibility than Vps8 (Shvarev et al., 2022). Importantly, Vps8 carries 282 additional amino acids compared to Vps41. Sequence alignment revealed that the main portion of these residues are inserted as 8 additional α-helices into the elbow region of Vps8 (Figure 3C).

The attachment of Vps8 to the core of the complex is similar to Vps3 and established through the long C-terminal α-helices and RING finger domains of the Vps8 and Vps18 subunits (Figures 1E, 2A,E,F). Reminiscent to the Vps11-Vps39 interface of HOPS (at the opposing end of the complex), the C-terminal helical bundle formed by Vps18 and Vps8 is followed by a pair of semi-symmetrically interacting RING finger domains (Figure 2F, upper panel). As shown for the Vps3-Vps11 interface, mutations in this interface (*vps18-1)* disassemble the entire complex (Figure S6G,H). Like Vps41 in HOPS, the C-terminus of Vps8 not only anchors the subunit to the core but also contributes to the attachment of the SNARE-binding module (Figure 2A-C). Distinctively, the binding interface between the core and the SNARE-binding module is larger in CORVET, which apparently increases stability. As in HOPS, Vps33 interacts with the RING finger domain of Vps18, and the N-terminal portion of Vps16 α-solenoid connects with the Vps8-Vps18 C-terminal long α-helical bundle (Figure 2A-C,E). Moreover, a hairpin protruding from the Vps8 RING finger domain is sandwiched in a hydrophobic groove between Vps33 and Vps18 (Figure 2B,C), clamping the SNARE-binding unit from both sides together with the C-terminus of Vps18 (Figure 2A,F upper panel). A similar interface likely exists in human HOPS, where the analogous Vps41 subunit also has a RING finger domain, which is lacking in yeast Vps41 (Kant et al., 2015; Radisky et al., 1997).

### Structural and functional implications of Vps8 interactions with Vps11 β-propeller

The hallmark of CORVET is the Vps8 subunit with its elbow shaped configuration. In addition to the C-terminal interaction with Vps18, and distinctive from HOPS, the α-solenoid of Vps8 has two additional interfaces with Vps11 (Figure 1D,E, 4A). The first interface is between two α-helices of Vps8 (residues 852-862 and 884-897) and a structured loop at the Vps11 β-propeller (residues 279-299) (Figure 4B), which stabilizes the upright section of the α-solenoid before the elbow. The second interface after the elbow connects three α-helices of Vps8 (residues 494-509, 534-550, 573-582) and the outward side of Vps11 β-propeller (residues 163-174) (Figure 4C).

**Figure 4.**
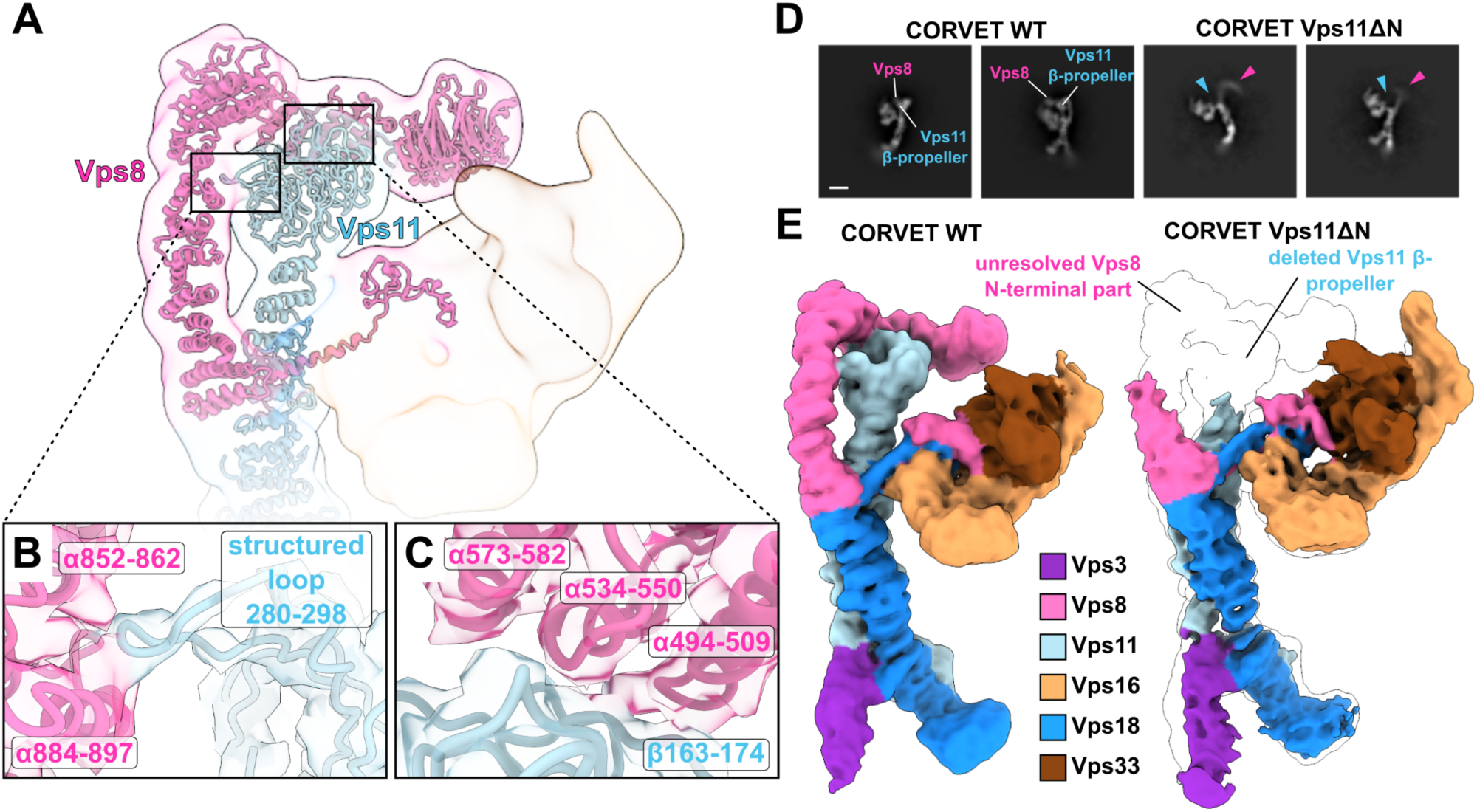
CORVET-specific Vps8-Vps11 N-terminal interface. **A**, Molecular models of CORVET Vps8 and Vps11 in ribbon representation fitted into the semi-transparent low-pass-filtered consensus cryo-EM map colored by subunits as in Figure 1. **B** and **C**, Zoomed-in views of the interface sites between Vps8 and Vps11 with interacting structural elements indicated. Associated semi-transparent cryo-EM densities are zoned around the molecular models and colored accordingly. **D**, Representative cryo-EM 2D class averages of CORVET wild-type and Vps111′N mutant. Structural differences in the regions of Vps8 and Vps11 N-termini are indicated. **E**, Comparison of CORVET wild-type and Vps111′N mutant cryo-EM densities colored by subunits as in Figure 1. The mutant cryo-EM density is overlayed on the wild-type density (transparent, black outline) highlighting the structural differences.

As these interfaces are distinct to CORVET, we investigated them closely. A mutant complex lacking the Vps11 β-propeller (CORVET Vps11βN, residues 1-335) could be purified like the wild-type complex (Figure S7A,C). Cryo-EM analysis indicates that this mutant not only lacks the β-propeller of Vps11, but shows poor resolution of N-terminal part of Vps8 (Figure 4D,E). We attribute this to increased flexibility in Vps8 due to the missing interface with Vps11.

We next explored if the corresponding mutation in CORVET affects its function. For HOPS, reconstituted tethering and fusion assays have been established (Ho and Stroupe, 2016; Lürick et al., 2017; Zick and Wickner, 2014), whereas comparable assays were missing so far for CORVET (Balderhaar et al., 2013; Lürick et al., 2017; Peplowska et al., 2007). We thus set up a similar fluorescence-based tethering assay for CORVET (Figure 5A) using Rab5(Vps21)-loaded liposomes (Langemeyer et al., 2020, 2018; Shvarev et al., 2022), including a screen for membrane conditions. Initially, we used vacuole mimicking lipid mix (VML) established for vacuole fusion assays (Zick et al., 2014) for the generation of liposomes (see Methods). Due to the unsaturated C18:2 lipids in the phospholipids, the corresponding liposomes have a more fluid membrane with packaging defects. This analysis revealed that CORVET requires relatively high PI3P (10 mol%) concentrations for efficient tethering (Figure 5B). To ask if membrane packaging defects contribute to efficient CORVET activity in tethering, we kept the VML composition for liposomes with the established PI3P concentration, but altered the acyl chain composition to palmitoyl (C16:0) oleolyl (C18:1) (PO) phospholipids (PO-VML), or used a more simple DL-phosphatidylcholine (PC) and DL-phosphatidylethanolamine (PE) mixture with the same PI3P concentration for liposomes (DLPC:DLPE). CORVET was only able to tether liposomes containing PI3P and the more fluid VML mixture, but none of the other liposomes (Figure 5C). This contrasts to HOPS, where the Rab7 GTPase Ypt7 is sufficient for tethering liposomes regardless of the composition (Figure 5C). We thus conclude that CORVET requires multiple factors – PI3P, Vps21 and membrane packaging defects – to bind membranes.

**Figure 5.**
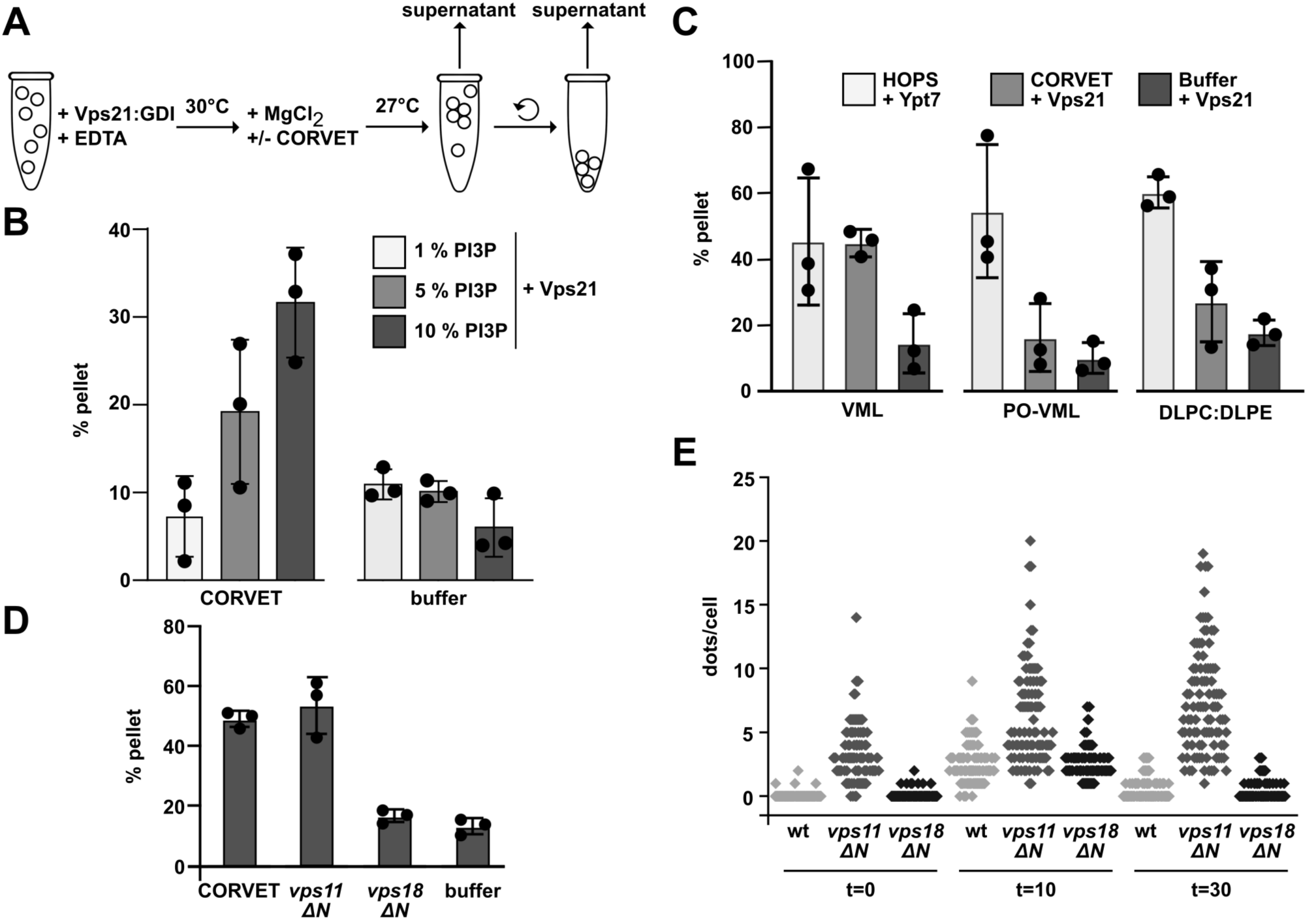
Functional analysis of CORVET-mediated membrane tethering. **A**, Scheme of the CORVET tethering assay (see Methods for details). **B**, Tethering assay with PI3P titration. Prenylated Vps21 was incorporated in fluorescently labelled liposomes containing 1, 5, or 10 mol % PI3P. Liposomes were incubated with purified CORVET complex or a buffer control. Tethering activity was determined as described in materials and methods. Dots indicate single measurements. Bars indicate standard deviation. Further tethering and statistics in Supplements. **C,** Tethering assay with DL– and PO-lipids. Liposomes containing 18:2/18:2 (DL-lipids) or 16:0/18:1 (PO-lipids) and SoyPI, PI3P, ergosterol and DAG or liposomes containing DLPC, DLPE and PI3P were fluorescently labelled. Prior to incubation with HOPS or CORVET liposomes were preloaded with prenylated Ypt7 or Vps21. Activity was measured according to materials and methods. Bars indicate standard deviation. Dots indicate single measurements. **D**, Tethering assay with CORVET carrying N-terminally truncated Vps11 and Vps18 subunits. Fluorescently labelled liposomes, decorated with prenylated Vps21, were incubated with wild-type or mutated CORVET variants. Tethering activity was measured as described in materials and methods. Bars indicate standard deviation. Dots indicate single measurements. **E**, Quantification of Mup1 uptake assay. Mup1 was GFP-tagged in wild-type, or cells expressing truncated *vps11ΔN* or *vps18ΔN*. Cells were grown in synthetic medium lacking methionine and imaged by fluorescence microscopy (t=0), before shifting to media containing methionine for 10 min (t=10) and 30 min (t=30). The number of GFP-positive dots per cell is plotted for each strain. Circles indicate single cells.

We then compared wild-type CORVET to mutants lacking either the Vps11 or Vps18 β-propeller (Vps18ΔN) in tethering assays. Surprisingly, the more flexible Vps11ΔN complex was still functional in tethering on membranes with high PI3P, whereas the Vps18ΔN was inactive (Figure 5D). The Vps18ΔN was fully assembled as it was purified as a hexamer like the wild type (Figure S7B), suggesting that the Vps18 β-propeller itself is critical for efficient tethering.

To determine if mutations in the β-propeller of Vps11 or Vps18 affect CORVET function, we used corresponding cells for possible endolysosomal defects (Figure S8). We initially analyzed vacuole morphology as CORVET deletion mutants have enlarged vacuoles (Banta et al., 1988; Cabrera et al., 2013; Raymond et al., 1992), but observed no difference to wild-type. However, cells expressing Vps11ΔN grew slower on plates containing endolysosomal stressors like ZnCl_2_ indicating possible defects (Figure S8B). To analyze early endocytic trafficking, we followed the transport of the GFP-tagged methionine transporter Mup1 from the plasma membrane via endosomal dots to the vacuole (Figures 5E, S8C). Whereas Mup1 was efficiently transported to the vacuole in wild-type and Vps18ΔN cells, it accumulated in GFP-positive dots over long periods in Vps11ΔN cells, suggesting impaired endosomal transport toward the vacuole in this mutant *in vivo*.

### Interaction of CORVET with membranes

To analyze how CORVET interacts with membranes, we first used *in silico* prediction of interactions of CORVET’s subunits Vps8 or Vps3 with the Rab5 GTPase Vps21. Our AlphaFold modeling (Evans et al., 2022) of the Vps3-Vps21 and Vps8-Vps21 interactions (Figure S9A-F) suggested binding sites on the peripheral areas of Vps8 and Vps3 α-solenoids, which would be accessible by membrane-bound Vps21 via its 10 nm long hypervariable domain (not shown in the prediction). The predicted interaction mode is similar to that in the HOPS complex (Shvarev et al., 2022), involving the same conserved residues in the Rab GTPases (Figure S9A-G), which are known to mediate Rab-effector binding (Barr, 2013; Müller and Goody, 2018).

To visualize CORVET on membranes, we incubated the complex with PI3P-containing liposomes loaded with Vps21-GTP and analyzed these by negative-stain microscopy. We observed CORVET particles coating the vesicle surfaces, mostly in an upright position (Figure S9H). Some particles also appeared flat on the membranes, possibly via Rab binding of both membrane-binding subunits in CORVET. The liposomes incubated without CORVET were indeed free of any decoration (Figure S9I). We conclude that CORVET is like HOPS on membranes preferentially upright prior to tethering.

## Discussion

In this study, we solved the structure of the crucial metazoan endolysosomal tethering complex CORVET. We show that CORVET and HOPS have modular architecture sharing an identical core with the attached SNARE-binding module and distinct complex-specific membrane-binding subunits. Importantly, we uncovered that CORVET differs strongly from HOPS in that it requires (i) PI3P, (ii) membrane packaging defects, and (iii) Rab5 (Vps21) for its efficient tethering function. Our data suggest that tethering complexes thus read out their membrane environment at several levels to gain organelle-specificity (Figure 6).

**Figure 6.**
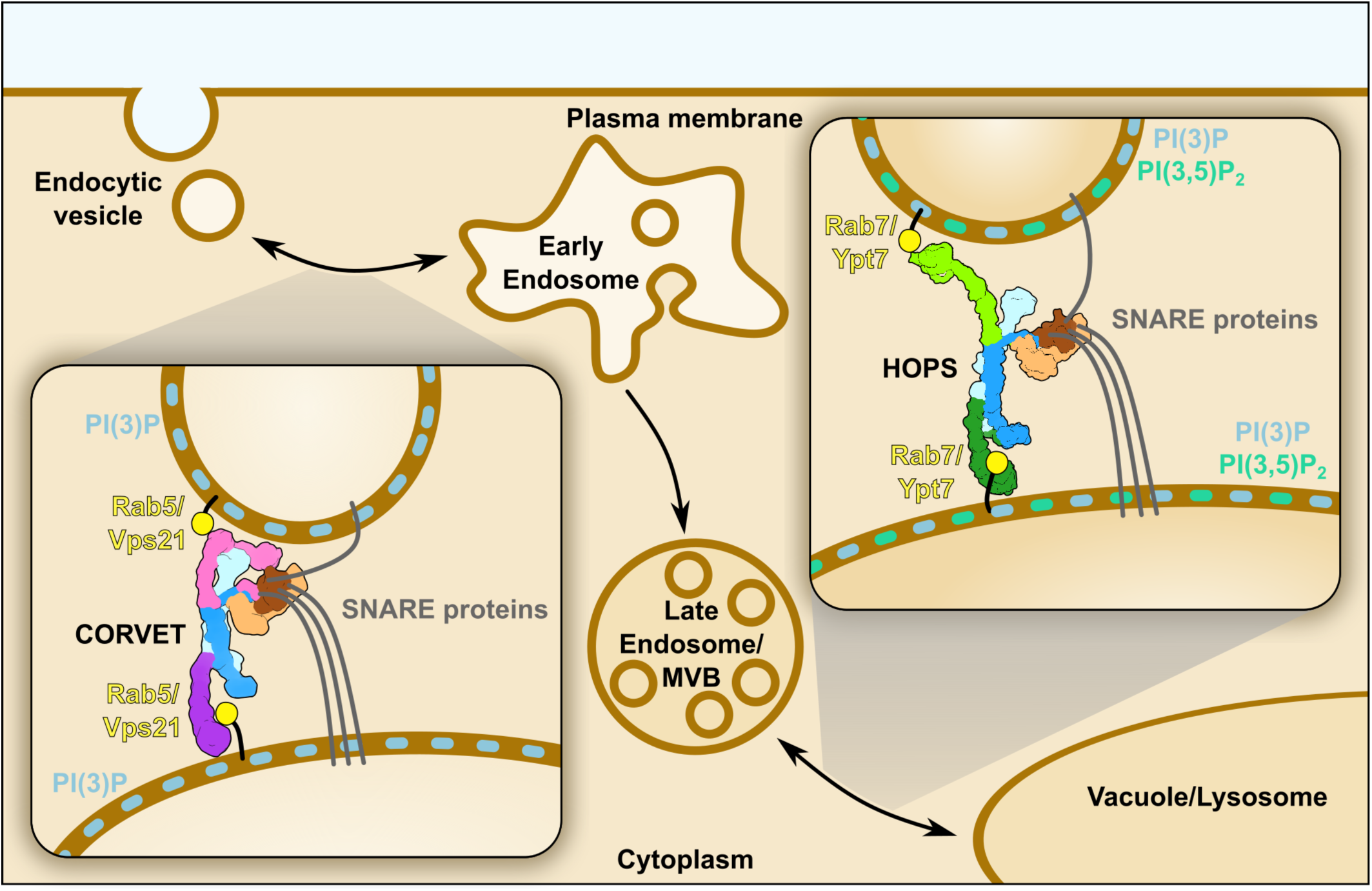
Mechanistic model of membrane tethering by CORVET and HOPS within the endolysosomal pathway. CORVET is responsible for early-endosomal fusion events and selectively relies on the Rab5 (Vps21) and PI3P presence in the membranes for function. In contrast, the lysosomal tether HOPS requires Rab7 (Ypt7) and, potentially, different lipid composition (e.g. PI(3,5)P_2_) to promote membrane fusion.

As in HOPS, subunit interactions in CORVET rely on the presence of C-terminal long α-helices and RING finger domains. As CORVET Vps3 misses a C-terminal RING finger domain, its association is weaker than Vps39 binding in HOPS, at least in solution (Ostrowicz et al., 2010; Peplowska et al., 2007). This is confirmed by our analysis of the Vps11-specific mutant alleles, which trigger CORVET disassembly *in vivo* (Figure S6C-F). It has been shown that RING finger domains of HOPS and CORVET subunits have ubiquitin ligase activity (Segala et al., 2019). As RING finger domains are structural elements of HOPS and CORVET, they may function as quality control elements to monitor the assembly of the complex. This could also explain, why an intermediate complex between HOPS and CORVET exists, which maintains the core part and can swap functional subunits such as Vps41/Vps8 or Vps39/Vps3, while the free subunit may fulfill auxiliary functions (Montoro et al., 2021; Peplowska et al., 2007). When Vps8 alone is overproduced, cells accumulate MVBs proximal to the vacuole (Markgraf et al., 2009), suggesting that Vps8 can constitute a minimal tether. Interestingly, a minimal CORVET complex, lacking Vps11 as a central subunit, was proposed in *Drosophila* (Lõrincz et al., 2016), which is, however, incompatible with the identified Vps11-Vps18 core of CORVET and HOPS. It is thus, not yet clear if miniCORVET is indeed a minimal CORVET or may have additional subunits.

One striking difference between CORVET and HOPS is the rigid apposition of Vps8 to the Vps11 β-propeller. This tight interaction can explain, why overproduced Vps8 in *Drosophila* can outcompete Vps41 and thus prevent HOPS assembly as it will likely sequester the available Vps11 pool (Lőrincz et al., 2019). The Vps8 apposition to the core and the Vps8-Vps11 contacts appear to be important for full CORVET activity in early endocytic transport *in vivo*. Interestingly, a reported point mutation in the α-solenoid of Vps8 might cause a destabilizing of the Vps8-Vps11 interaction (Plemel et al., 2011). We demonstrated this by our structural (Figure 4) and functional (Figure 5) analyses of CORVET Vps11ΔN. Indeed, a deletion of the β-propeller domain of core subunits such as Vps11 and Vps18 will affect both CORVET and HOPS *in vivo*. In this regard, we were surprised that the CORVET Vps18ΔN mutant, in contrast to Vps11ΔN, was largely inactive in the tethering assay (Figure 5D), whereas the comparable HOPS complex was functional (Shvarev et al., 2022). This suggests that Vps18 either contributes to membrane recognition by Vps3 within CORVET or to the structural integrity of the complex. In either case, we can only speculate, how membranes are decoded by CORVET and how the β-propeller domains of Vps11 and Vps18 contribute. One idea is that the β-propeller of core Vps11 and Vps18 bind Rab5 GTPases together with the Rab5-GTP specific subunits Vps3 and Vps8, thus optimally positioning them for direct PI3P and membrane binding.

We show that CORVET depends on multiple membrane cues, whereas HOPS can tether any membrane carrying the Rab GTPase Ypt7. For membrane binding, CORVET may thus require specific lipids and membrane packaging defects, either through high curvature or due to the membrane composition (Bigay and Antonny, 2012; Holthuis and Menon, 2014), in addition to Rab5, which is needed for CORVET localization *in vivo* (Cabrera et al., 2013). In line with this, predicted Rab binding sites of CORVET and HOPS are largely conserved and located at the distal exposed ends of the complexes. However, Rab5 also binds and stimulates the Vps34 PI-3-kinase complex, which establishes a suitable membrane environment required for CORVET recognition (Christoforidis et al., 1999; Tremel et al., 2021). Consequently, loss of Rab5 (or its homolog in yeast) will possibly affect both the CORVET binding and the specific membrane environment.

Currently, we have no clear understanding, how tethering complexes function on membranes to promote tethering and fusion. Our analyses of HOPS suggests that the complex is positioned upright on membranes when bound to the Rab7-like Ypt7 protein (Füllbrunn et al., 2021). Here, we used liposomes and find CORVET also mostly upright on membranes (Figure S9H). This suggests that direct interactions with the lipid bilayer strongly contribute to the positioning of CORVET, making it ready for tethering and subsequent promotion of SNARE-mediated fusion, while Rab GTPases may further stabilize the complex.

Our analysis of CORVET reveals that its core architecture largely resembles HOPS, yet the orientation and structure of the Rab-specific subunits and their decoding of membranes strongly differs between both complexes. Nevertheless, though interchangeable assemblies of CORVET and HOPS are possible, the four shared central subunits, Vps11, 18, 33 and 16, are found in all species (Klinger et al., 2013). We are thus convinced that the overall arrangement and function of HOPS and CORVET as a modular membrane fusion machinery is evolutionary conserved. Future studies will take advantage of our structural insights to determine, how these complexes promote SNARE-mediated membrane fusion.

## Acknowledgments

The authors thank Dovile Januliene for help with cryo-EM experiments, Kilian Schnelle for help with computing and data processing, and all members of the Ungermann and Moeller lab for feedback. This work was supported by a grant of the DFG (UN111/5-6; MO2752/3-6), the SFB 944 and SFB 1557 (C.U.; F.F., A.M.), the DFG INST190/196-1 FUGG (A.M.) and the BMBF/DLR 01ED2010 (A.M.).

## Author contributions

D.S., C.K. and N.S. conceived and designed all experiments together with A.M. and C.U.. D.S. conducted all cryo-EM analyses. C.K. and N.S. did all biochemical and cell biology experiments. F.F. performed mass spectrometry analysis with support of S.W. The manuscript was written by D.S., C.U. and A.M. with contributions from all authors.

## Data and materials availability

The cryo-EM density maps and corresponding atomic models reported in this study have been deposited in the Electron Microscopy Data Bank and Protein Data Bank with the accession codes EMD-xxx and PDB-xxx, respectively.

## METHODS

### Yeast strains

Yeast strains and oligonucleotides used in this study are listed in Table S1 and Table S2 respectively. In general, CORVET subunits were expressed under the control of the GAL1 promoter according to the standard protocol (Janke et al., 2004). For subunit truncation (Vps11 or Vps18) of the N-terminal part, the GAL1 promotor was inserted into the genome at the respective position. The 3x-FLAG Tag was attached to the CORVET subunit Vps8, except for the Vps18ΔN mutant.

### Protein expression and purification from *Escherichia coli*

Rab GTPases for tethering assays were expressed in *Escherichia coli* BL21 (DE3) Rosetta cells. Cultures were grown in Luria broth (LB) medium complemented with 35 µg/ml kanamycin or 100 µg/ml ampicillin and 30 µg/ml chloramphenicol. Cultures were induced with 0.5 mM Isopropyl-β-D-thiogalactoside (IPTG) and incubated overnight at 16°C. Cells were harvested by centrifugation (4800*×g*, 10 min, 4°C) and resuspended in buffer (150 mM NaCl, 50 mM HEPES/NaOH, pH 7.4, 10 % glycerol, 1 mM PMSF, and 0.5-fold protease inhibitor mixture [PIC]) before lysed in a Microfluidizer (Microfluidics Inc). Crude lysate was centrifugated at 25,000*×g*, 30 min, 4°C. Supernatants were incubated with glutathione Sepharose (GSH) fast flow beads (GE Healthcare) for GST-tagged proteins or nickel– nitriloacetic acid (Ni-NTA) agarose (Qiagen) for His-tagged proteins (2 hr, 4°C). Proteins were eluted with buffer (150 mM NaCl, 50 mM HEPES/NaOH, pH 7.4, 10 % glycerol) containing either 25 mM glutathione or 300 mM imidazole. Buffer was exchanged via a PD10 column (GE Healthcare). For tag cleavage, TEV or PreScission protease was added after washing and incubated overnight. Proteins were stored at −80°C.

### Purification of 3xFLAG-tagged CORVET complex variants

CORVET tethering complex variants were purified according to the standard FLAG-purification protocol as previously described (Shvarev et al., 2022) with minor changes. Two litres medium (yeast peptone (YP), containing 2 % galactose (v/v)) were inoculated with 6 ml of an overnight preculture. Cultures were grown for 24 hr and harvested by centrifugation (4800*xg*, 10 min, 4°C). Cell pellets were washed with cold CORVET purification buffer (CPB, 300 mM NaCl, 20 mM HEPES/NaOH, pH 7.4, 1.5 mM MgCl_2_, and 10% (v/v) glycerol). Pellets were resuspended in CORVET lysis buffer (CLB, CPB supplemented with 1 mM phenylmethylsulfonylfluoride (PMSF), 1× FY protease inhibitor mix (Serva) and 1 mM dithiothreitol (AppliChem GmbH)). Cell suspension was dropwise frozen in liquid nitrogen before being lysed in a freezer mill cooled with liquid nitrogen (SPEX SamplePrep LLC). For purification, the powder was thawed on ice and resuspended in CLB, followed by two steps of centrifugation at 5000 and 15,000*×g* at 4°C for 10 and 20 min. The supernatant was combined with anti-FLAG M2 affinity gel (Sigma-Aldrich) and placed on a nutator for 45 min at 4°C. Beads were centrifugated (500*×g*, 1 min, 4°C) and transferred to a 2.5 ml MoBiCol column (MoBiTec). Samples were washed with 25 ml CPB before FLAG-peptide was added, followed by incubation on a turning wheel for 40 min at 4 °C. The eluate was concentrated in a Vivaspin 100kDa MWCO concentrator (Sartorius), which was incubated with CPB containing 1 % TX-100. Concentrated sample was applied to a Superose 6 Increase 15/150 column (Cytiva) for size exclusion chromatography (SEC). Fractions were eluted in 50 μl using ÄKTA go purification system (Cytiva). Peak fraction was used for further analysis.

### Mass photometry analysis

Mass photometry experiments were performed using a Refeyn TwoMP (Refeyn Ltd). Data were acquired using AcquireMP software and analyzed using DiscoverMP (both Refeyn Ltd). High Precision Cover Glasses (Marienfeld) were used for sample analysis. Perforated silicone gaskets were placed on the coverslips to form wells for every sample to be measured. Samples were evaluated at a final concentration of 10 nM in a total volume of 20 μl in the buffer used for SEC. Calibration was performed using β-amylase (Carl Roth).

### ALFA Pulldowns for mass spectrometry

Vps8-ALFA pulldowns were performed in the same way as previously described (Shvarev et al., 2022). One liter YP medium containing 2% glucose (v/v) was inoculated with an overnight preculture. Cells were grown to OD600 1 at 26°C, followed by 1 hr heat shock at 38°C. Cells were harvested by centrifugation (4800*×g* for 10 min at 4°C). Pellets were washed with cold Pulldown buffer (PB, 150 mM KAc, 20 mM HEPES/NaOH, pH 7.4, 5% (v/v) glycerol and 25 mM CHAPS). Cells were resuspended in a 1:1 ratio (w/v) in PB supplemented with Complete Protease Inhibitor Cocktail (Roche) and afterward dropwise frozen in liquid nitrogen before lysed in 6875D LARGE FREEZER/MILL (SPEX SamplePrep LLC). Powder was thawed on ice and resuspended in PB, followed by two centrifugation steps at 5000 and 15,000×g at 4°C for 10 and 20 min. Supernatant was added to 12.5 µl prewashed ALFA Selector ST beads (2500*×g*, 2 min, 4°C) (NanoTag Biotechnologies) and incubated for 15 min at 4°C while rotating on a turning wheel. Afterwards, beads were washed two times in PB and four times in PB without CHAPS. Samples were digested using PreOmics sample kit (iST Kit, preomics) and analyzed in Q ExativePlus mass spectrometer (Thermo Fisher Scientific).

### Negative stain analysis

Samples of wild-type CORVET, mutant CORVET variants, and CORVET incubated with liposomes were examined by negative-stain electron microscopy. 3 μl of sample at a protein concentration of approximately 0.05 mg/ml was applied onto freshly glow-discharged carbon-coated copper grids with plastic support, blotted from the side and stained using 2% (w/v) uranyl formate solution as previously described (Januliene and Moeller, 2021). Negative-stain micrographs were collected manually on a JEM-2100Plus transmission electron microscope (JEOL) operated at 200 kV and equipped with a XAROSA CMOS 20-megapixel camera (EMSIS) at a nominal magnification of 30,000 (3.12 Å per pixel). The data was analyzed using ImageJ (Schneider et al., 2012) and cryoSPARC (Punjani et al., 2017).

### Cryo-EM sample preparation and data acquisition

For cryo-EM, 3 μL of freshly purified CORVET samples at a protein concentration of approximately 0.8 mg/ml were applied to glow-discharged C-Flat grids (R1.2/1.3 3Cu-50) (EMS) and immediately plunge frozen in liquid ethane using a Vitrobot Mark IV (Thermo Fisher Scientific) with the environmental chamber set at 100% humidity and 4°C.

Micrographs were collected automatically with EPU (Thermo Fisher Scientific), using a Glacios cryo-transmission electron microscope (Thermo Fisher Scientific) operated at 200 kV and equipped with a Selectris energy filter and a Falcon 4 detector (both Thermo Fisher Scientific). Data were recorded in Electron Event Representation (EER) mode at a nominal magnification of 130,000 (0.924 Å per pixel) in the defocus range of −0.8 to −1.8 μm with an exposure time of 7.50 s resulting in a total electron dose of approximately 50 e^−^ Å^−2^.

### Cryo-EM image processing

All cryo-EM data were preprocessed in cryoSPARC Live, and further processing was performed in cryoSPARC v3 and v4 (Punjani et al., 2017) (Figure S2). For all collected movies, motion correction (EER upsampling factor 1, EER number of fractions 40) and contrast transfer function (CTF) estimation were performed using cryoSPARC Live implementations. After micrograph curation, 33882 micrographs were included for further data analysis.

For the determination of the wild-type structure, all collected datasets were preprocessed in a similar manner and combined at the final steps of image processing (Figure S2). Briefly, particles were selected by the template picker implemented in CryoSPARC Live, using well-defined 2D classes obtained from preliminary CORVET datasets as templates, and additional particle picking was performed using the Topaz wrapper (Bepler et al., 2019). For particle picking, micrographs from sessions 1-5 were combined, while the micrographs from sessions 6 and 7 were used separately. After particle duplicate removal and particle extraction in a box size of 882 pixels (Fourier-cropped to 128 pixels, resulting in a pixel size of 6.37 Å per pixel), two rounds of 2D classification were performed for all particle stacks obtained from each of the picking jobs to eliminate bad picks (Figure S2). After 2D classifications, duplicate particles were removed again and a round of ab-initio reconstruction with 5 classes followed by heterogeneous refinement was performed for each stack of particles resulted from sessions 1-5, session 6, and session 7. Each of these three heterogeneous refinement jobs produced one best class revealing all of the subunits of CORVET in the map. Such a class resulted from session 7 was further refined individually using NU refinement (Punjani et al., 2020) and particles from it (158046 particles) were afterwards combined with the particles from the best classes of the heterogeneous refinement jobs from sessions 1-5 (93661 particles) and 6 (216338 particles). A round of heterogeneous refinement with three classes was performed using these combined particles in a box size of 882 pixels (Fourier-cropped to 224 pixels, pixel size 3.64 Å per pixel), which resulted in two good classes reaching resolution of 7.4 Å (Gold Standard Fourier Shell Correlation (GSFSC) value of 0.143). These two classes (174263 and 198464 particles) were combined and further refined using NU refinement in a box size of 882 pixels (Fourier-cropped to 400 pixels, pixel size 2.04 Å per pixel). The obtained reconstruction resolved to 4.6 Å (GSFSC = 0.143) was further classified using a round of heterogeneous refinement with 5 classes. The best class that reached resolution of 6.7 Å (GSFSC = 0.143) was used for a NU refinement providing a consensus map (219391 particles) at 4.6 Å resolution (GSFSC = 0.143). This consensus map was further used for local refinements of distinct areas of the structure. For local refinements, five masks were generated using UCSF ChimeraX (Pettersen et al., 2020) and particles were again re-extracted for some of the parts of the structure in a box size of 882 pixels (Fourier-cropped to 512 pixels, pixel size 1.59 Å per pixel) (Figure S2). A composite map was generated from the local refinement maps using the “volume maximum” command in UCSF ChimeraX.

All maps were subjected to unsupervised B-factor sharpening within cryoSPARC. No symmetry was applied during processing. The quality of the consensus and local refinement maps is shown in Figures S3 and S4. All GSFSC curves and angular distribution plots were generated within cryoSPARC (Figure S3). The local resolutions of the consensus and local refinement maps (Figure S3) were estimated in cryoSPARC and analyzed in UCSF ChimeraX. Dataset statistics are provided in Table S3. The cryo-EM analysis of the CORVET Vps11ΔN mutant was performed similarly to the wild type (Figure S7D).

### Model building and refinement

The structures of Vps11, Vps16, Vps18, and Vps33 from the structure of the HOPS complex (PDB: 7ZU0) and the AlphaFold structure predictions of Vps8 and Vps3 (Uniprot: P39702/ P23643 respectively) were manually fitted into the consensus and local-refinement maps using the “Fit in Map” tool in UCSF ChimeraX and used as a starting model. Most of the structure was modelled as poly-alanine sequences using the PDB Tools job in PHENIX (Liebschner et al., 2019), except for the regions of the structure with the highest resolution obtained (SNARE-binding module, Vps8 α-solenoid, N-terminal half of Vps11), in addition, several protein fragments without well resolved corresponding EM densities were removed from the model (including the N-terminal half of Vps3). The models of the CORVET subunits were then manually adjusted and refined in Coot (Emsley et al., 2010) and combined into a single model of the full complex. Subsequently, iterative rounds of real space refinement (Afonine et al., 2018) of the model against the composite map of CORVET in PHENIX were performed, followed by manual adjustments in Coot. Model validation was done using MolProbity (Williams et al., 2018) in PHENIX. Models and maps were visualized, and figures were prepared in UCSF ChimeraX and Inkscape. Model refinement and validation statistics are provided in Table S3.

### Liposome preparation

For the preparation of liposomes, the following lipids were used. 1,2-dilinoleoyl-sn-glycero-3-phosphocholine (18:2 PC), 1-palmitoyl-2-oleoyl-glycero-3-phosphocholine (POPC) 1,2-dilinoleoyl-sn-glycero-3-phosphoethanolamine (18:2 PE), 1-palmitoyl-2-oleoyl-sn-glycero-3-phosphoethanolamine (POPE), L-α-phosphatidylinositol (SoyPI), 1,2-dilinoleoyl-sn-glycero-3-phospho-L-serine (18:2 PS), 1-palmitoyl-2-oleoyl-sn-glycero-3-phospho-L-serine (POPS), 1,2-dilinoleoyl-sn-glycero-3-phosphate (18:2 PA), 1-palmitoyl-2-oleoyl-sn-glycero-3-phosphate (POPA), 1,2-dipalmitoyl-sn-glycero-3-(cytidine diphosphate) (DAG) were purchased from Avanti Polar Lipids (Alabama, USA). Phosphatidylinositol 3-phosphate (PI3P) was purchased from Echelon Biosciences Inc (Utah, USA). ATTO488-1,2-dipalmitoyl-*sn*-glycero-3-phosphoethanolamine (ATTO488) was obtained from ATTO-TEC GmbH (Siegen, Germany). Ergosterol was purchased from Sigma Aldrich.

For tethering assays and negative stain, lipid films, containing 37.6 or 42.6 mol% 18:2 PC/ POPC, 18 mol% 18:2 PE/ POPE, 18 mol% SoyPI, 10 or 5 mol % PI(3)P, 4.4 mol% 18:2 PS/ POPS, 2 mol% 18:2 PA/ POPA, 8 mol% Ergosterol, 1 mol% DAG and 1 mol% ATTO488, were evaporated by a SpeedVac (CHRIST, Germany). Lipid films prepared for the negative stain EM analysis did not contain dye, and the 18:2 PC content was raised accordingly. Lipid films were resuspended in buffer (25 mM HEPES/NaOH, pH 7.4, 135 mM NaCl). Unilamellar vesicles were obtained through seven freeze and thaw cycles in liquid nitrogen. Liposomes were extruded through polycarbonate filters to 100 nm for tethering assays and 30 nm for negative stain EM analysis (400 nm, 200 nm, 100 nm, 50 nm and 30 nm pore size) using a hand extruder (Avanti Polar Lipids, Inc.).

### Tethering assay

CORVET-mediated tethering assays were done as before (Lürick et al., 2017). ATTO488 labelled liposomes were prepared and loaded with prenylated Vps21 or Ypt7 (Langemeyer et al., 2018). 100 nmole liposomes were incubated with 100 pmole pVps21:GDI/Ypt7:GDI in the presence of 1 mM GTP and 20 mM EDTA for 30 min at 30° C, before the addition of 25 mM MgCl_2_. 100 nM CORVET/HOPS or buffer (HEPES/NaOH, pH 7.4, 300 mM NaCl, 1.5 mM MgCl_2_) were incubated with 0.17 mM Rab-loaded liposomes for 10 min at 27° C. Liposomes were sedimented by centrifugation at 2,500 *xg* for 5 min. The tethered liposomes in the pellet fraction were determined by comparing the ATTO488 fluorescent signal of the supernatant before and after centrifugation using a SpectraMax M3 fluorescence plate reader (Molecular Devices). Statistical analyses were performed with GraphPad Prism 9.

### Imaging of yeast cells using fluorescence microscopy

Cells were grown to the exponential phase in synthetic complete medium supplemented with all amino acids (SDC), or lacking methionine (SDS-met) [0.675% (w/v) yeast nitrogen base without amino acids, 2.0% (w/v) glucose, 0.075% (w/v) CSM (MPBiomedicals)]. Cells were imaged in SDC or SDC-met using the Zeiss Axioscope 5, with a PLAN-Apochromat 100x/1.40 Oil DIC M27 objective and an axiocam 702 mono (1.0x) camera.

All images were processed and quantified using ImageJ NIH, Bethesda, MD. Statistical analyses were performed with Origin Pro 9 software. Vacuoles were stained using 30 µM FM4-64 (Molecular Probes Inc., Eugene, OR) for 30 min. Cells were washed twice with medium, prior to 1 h incubation at 30 °C (Vida and Emr, 1995).

