## Supplementary Material for "Structure of the endosomal CORVET tethering complex"

#### Supplementary Figures

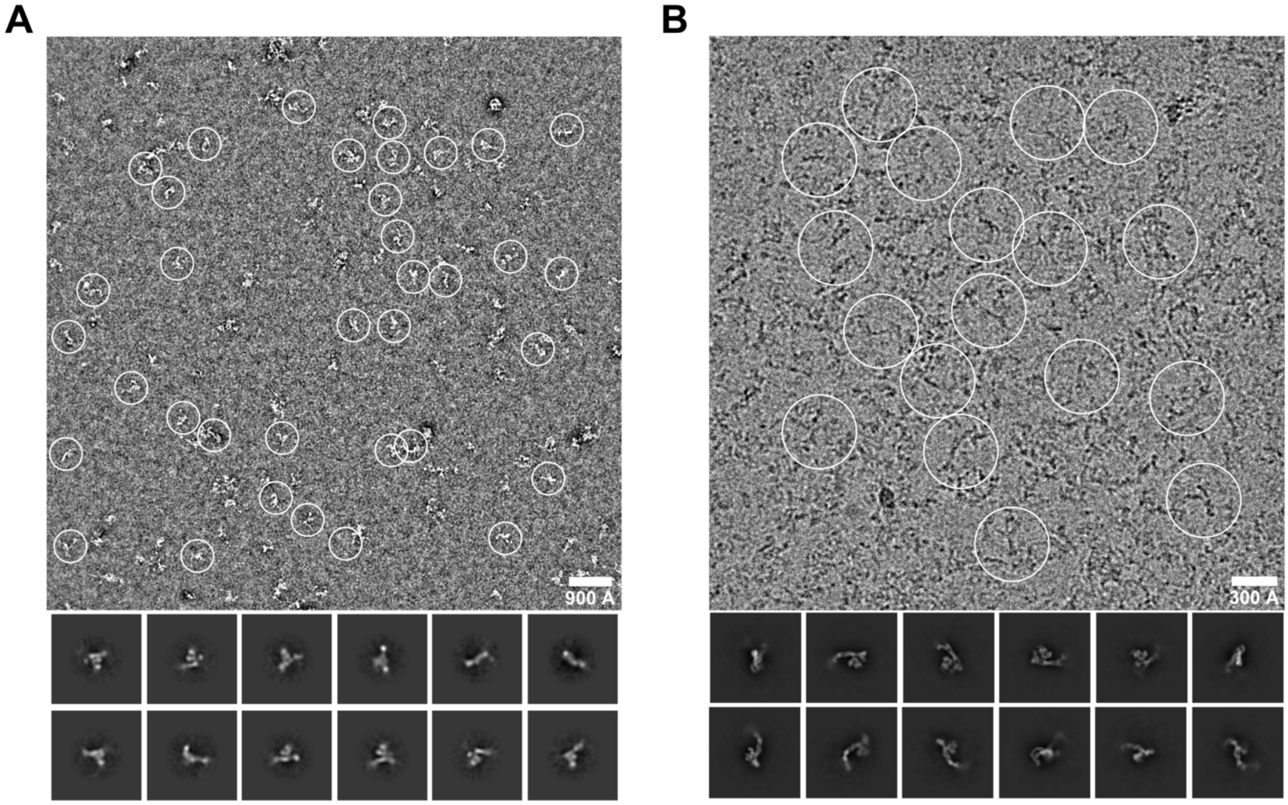


**Figure S1. Single particle analysis of wild-type CORVET.** Representative negative stain **(A)** and cryo-EM **(B)** micrographs and 2D class averages (bottom). Box size used for particle extraction: 799 Å (negative stain), 815 Å (cryo-EM).

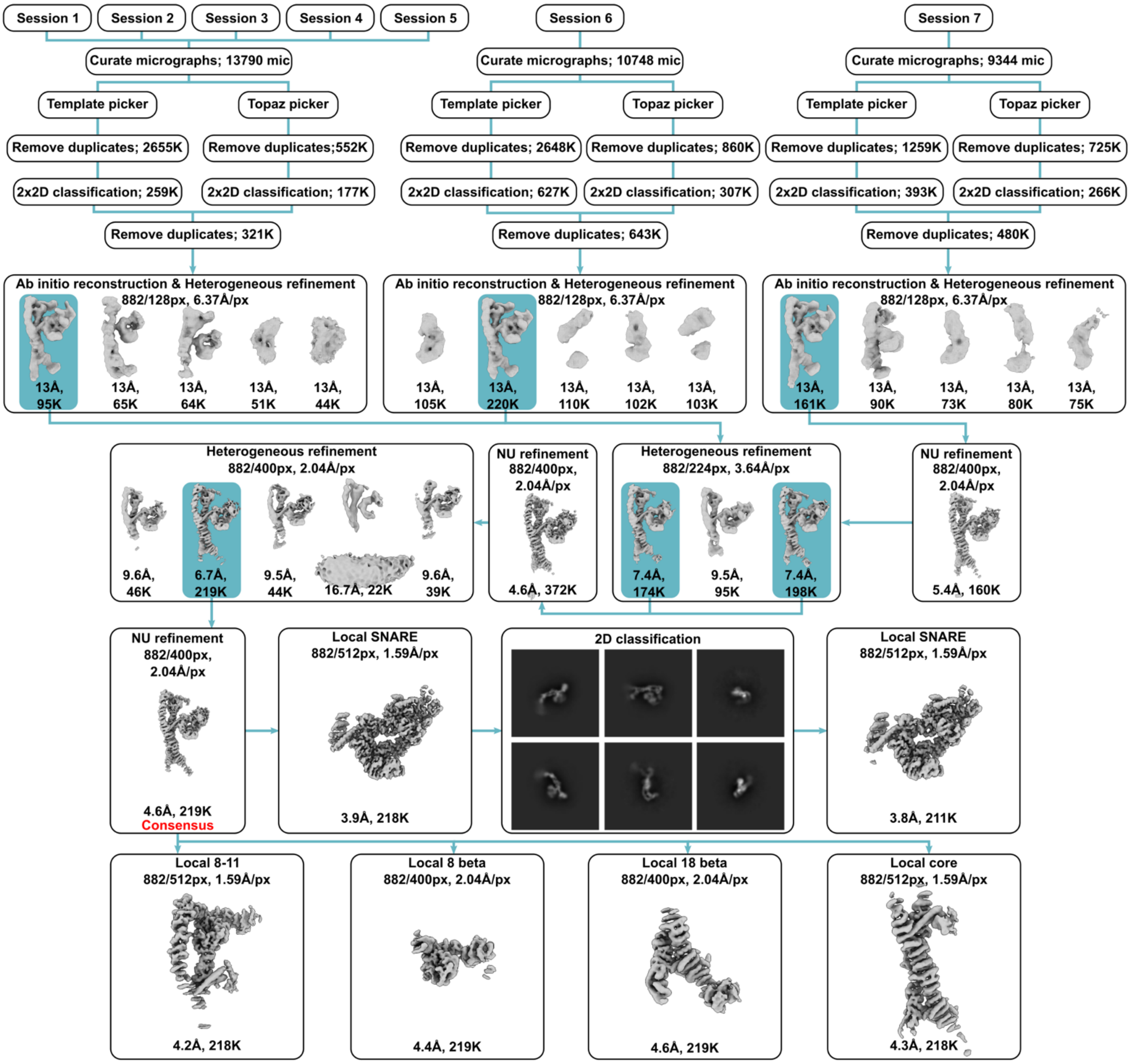


**Figure S2**. **Cryo-EM data processing workflow of wild-type CORVET.** The main steps of the data processing pipeline performed in cryoSPARC are shown. Box sizes (full/cropped) in pixels, resulting pixel size, number of particles, and resolution achieved are shown for each map shown.

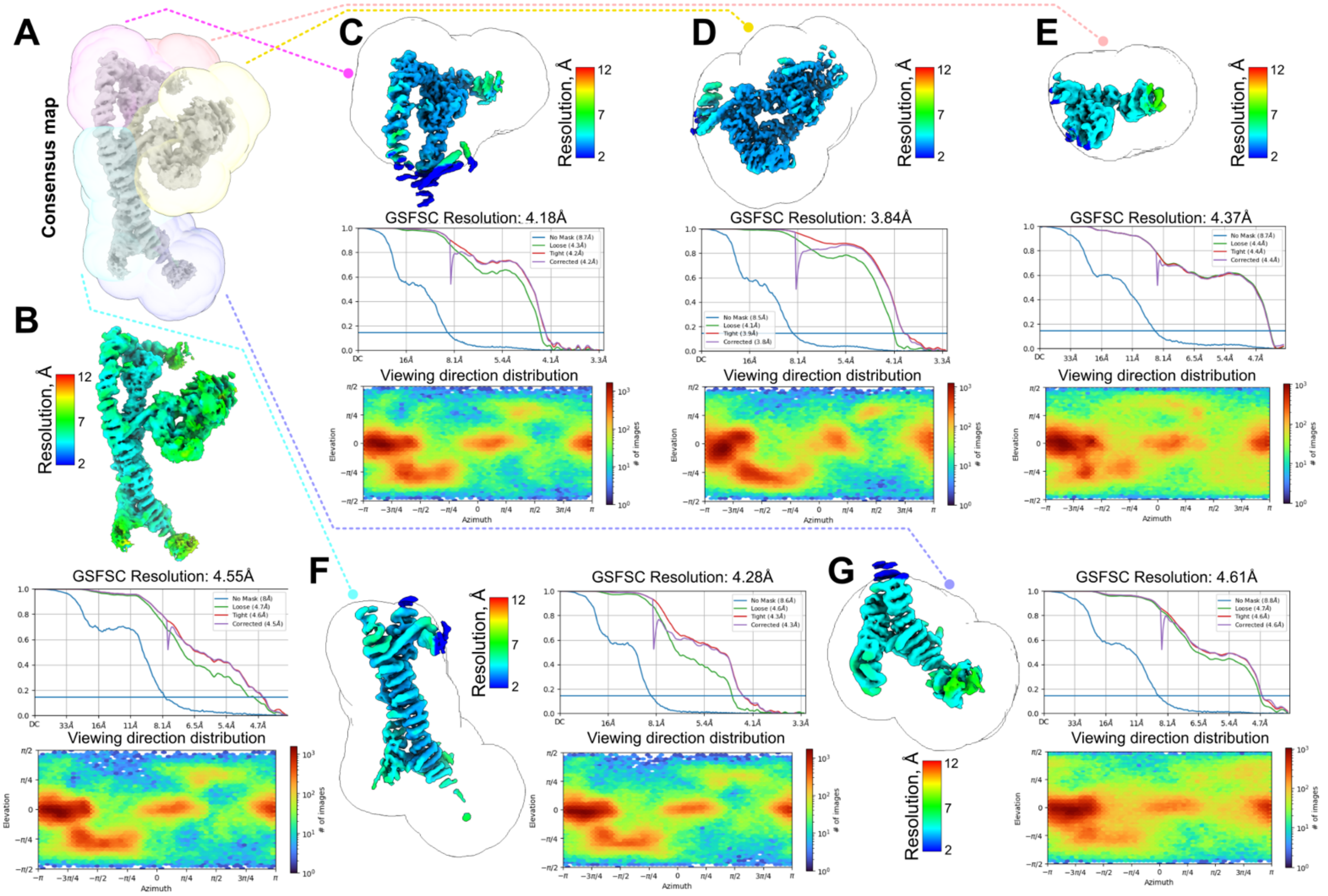


**Figure S3**. **Local refinement approach and cryo-EM data validation. A,** Masks (semi-transparent) used for local refinement of the consensus map fragments. **B-G,** Local resolution estimation, FSC curve, and angular distribution heatmap plot generated in cryoSPARC for the consensus map **(B)** and final local refinement maps **(C-G)**.

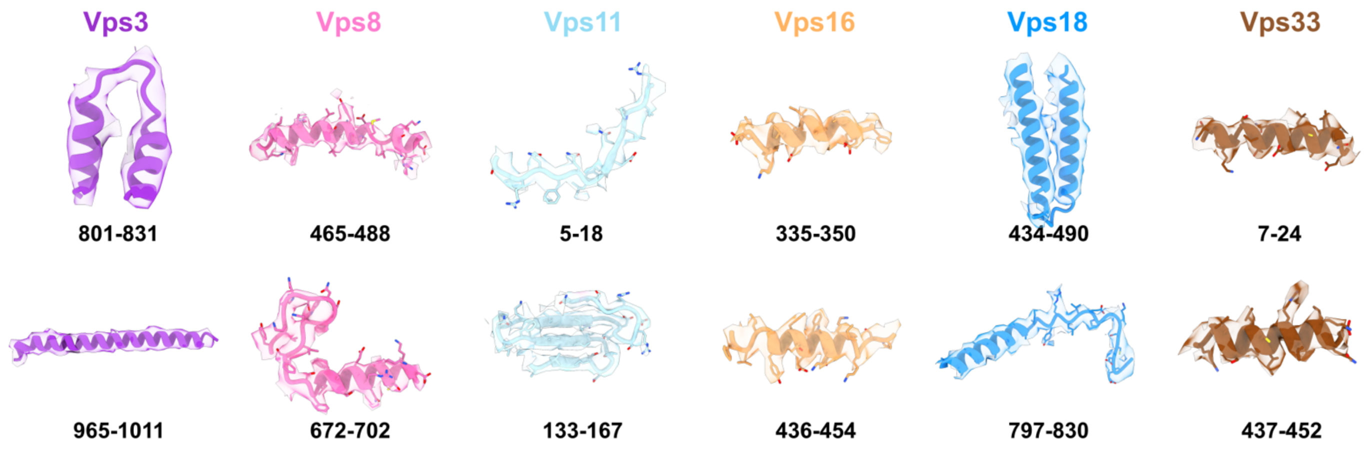


**Figure S4**. **Cryo-EM quality.** Model/map fit of selected areas within each subunit of CORVET.

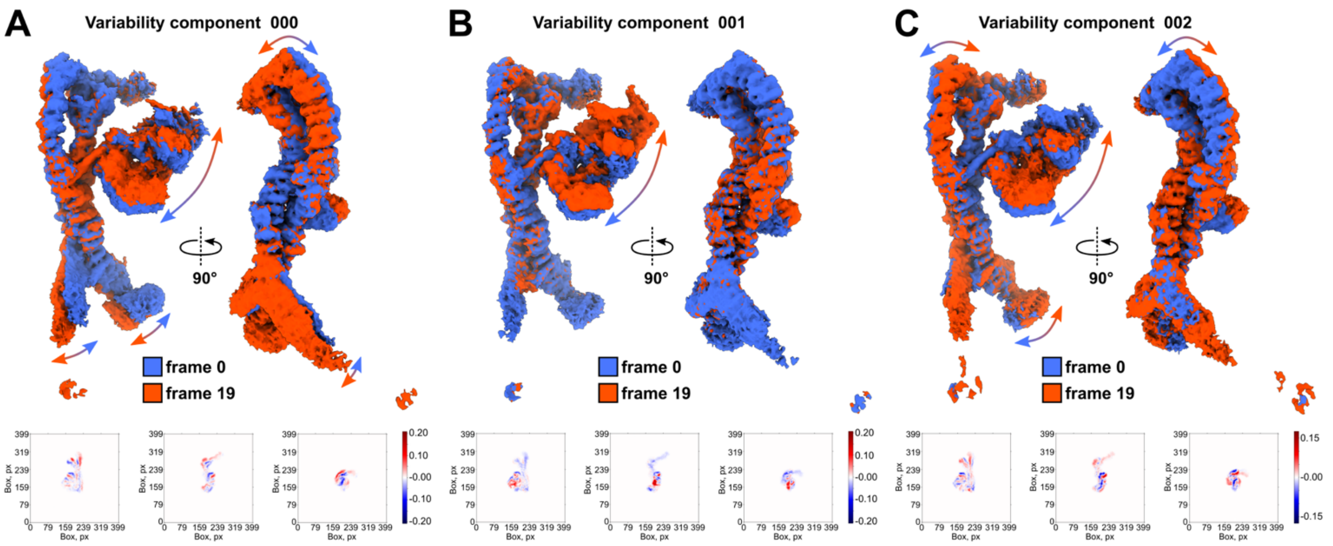


**Figure S5**. **3D variability analysis of CORVET. A-C,** 3D density maps (top) and map 2D projections (bottom) at negative (blue) and positive (red) positions along each variability component are shown.

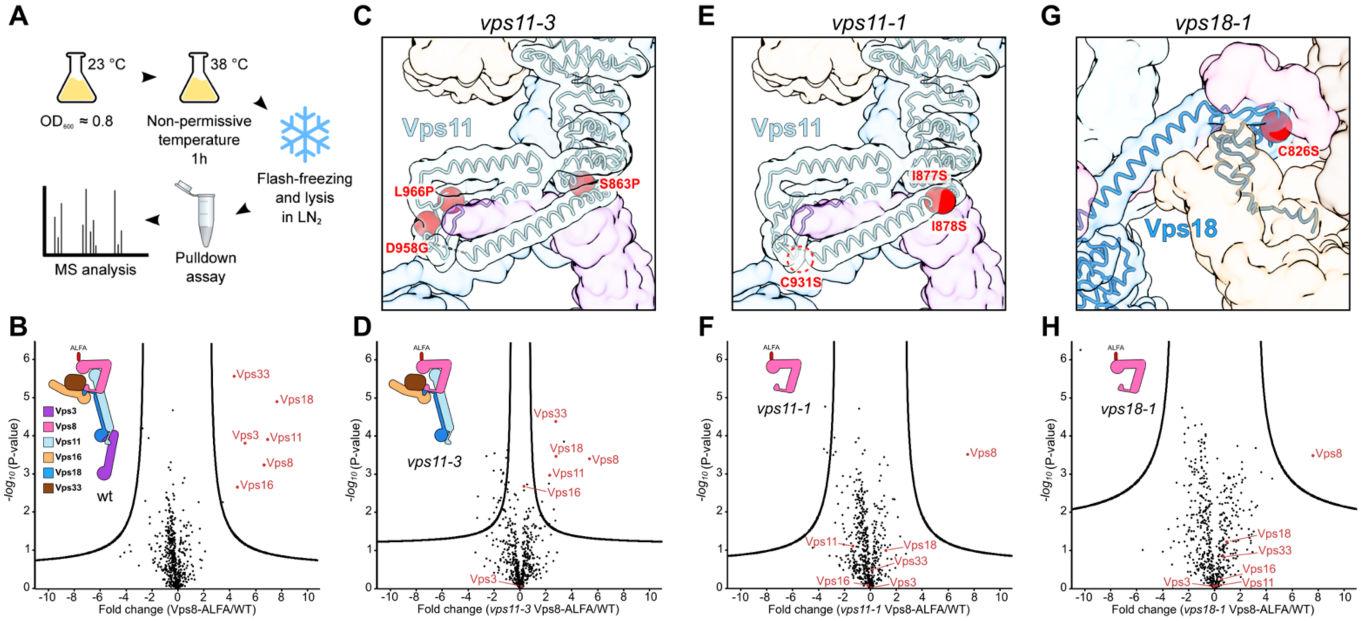


**Figure S6**. **Analysis of RING finger domains role in the structural integrity of CORVET. A,** Scheme of the experiment setup. **B,D,F,H**, Mass spectrometry analysis of Vps8 purified via the ALFA tag from the indicated strains and enriched proteins (red dots). Results of purification from wt (**B**), *vps11-3* (**D**), *vps11-1* (**F**), and *vps18-1* (**H**) cells. **C,E,G**, Positions of the mutated residues (red spheres) in the respective mutants of the core subunits Vps11 or Vps18 (shown in ribbon representation) in the context of the CORVET structure (semi-transparent envelope).

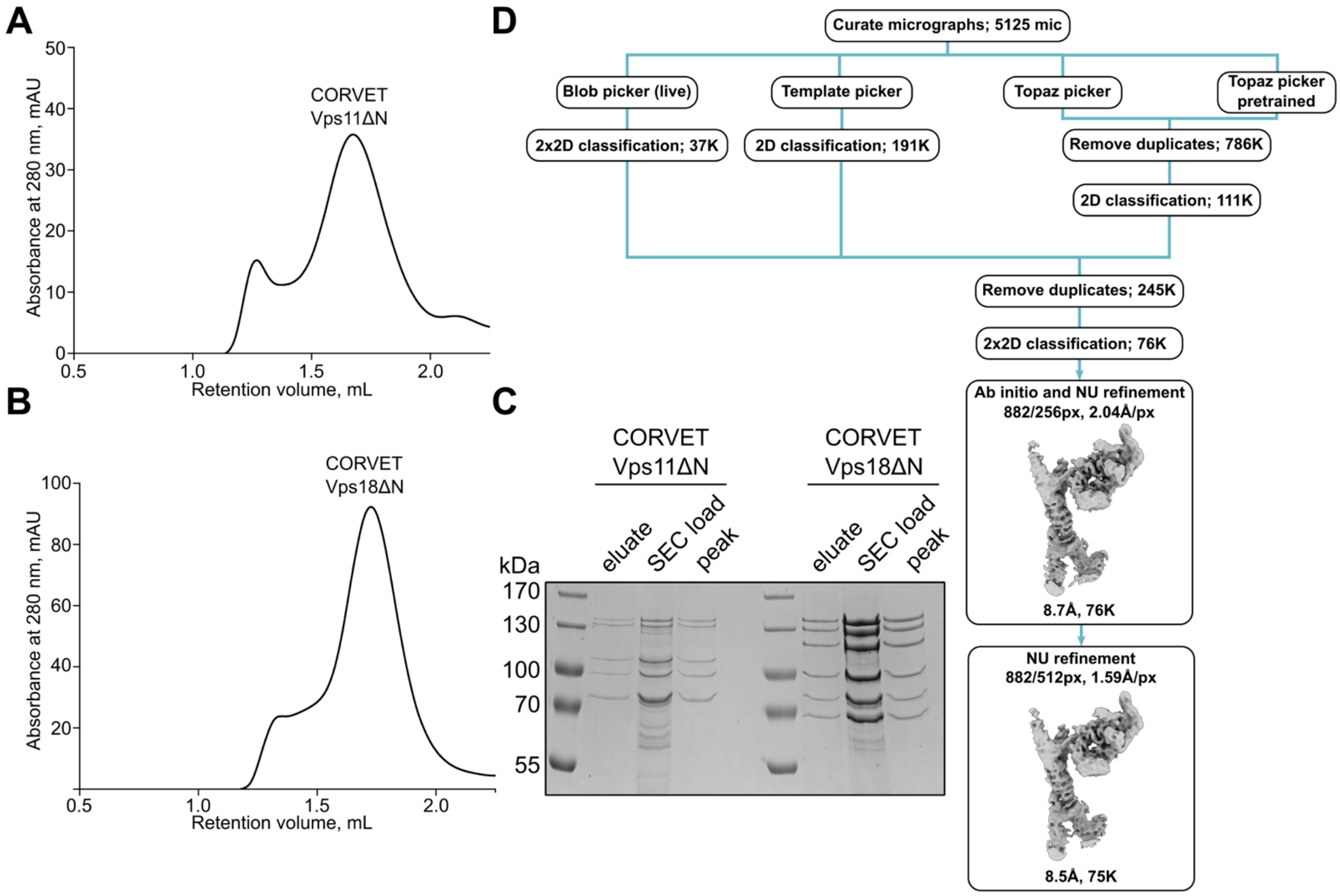


**Figure S7**. **Purification, biochemical analysis and cryo-EM workflow of the CORVET Vps11ΔN mutant complex. A,** Size exclusion chromatography (SEC) of the affinity-purified CORVET Vps11ΔN. **B**, Size exclusion chromatography (SEC) of the affinity-purified CORVET Vps18ΔN. **C**, SDS-PAGE analysis of purified CORVET Vps11ΔN and Vps18ΔN mutant complexes. Protein samples from affinity purification (eluate) and before and after SEC are shown. **D**, The main steps of the data processing pipeline of the cryo-EM CORVET Vps11ΔN sample performed in cryoSPARC are shown. Box sizes (full/cropped) in pixels, resulting pixel size, number of particles, and resolution achieved are shown for each map shown.

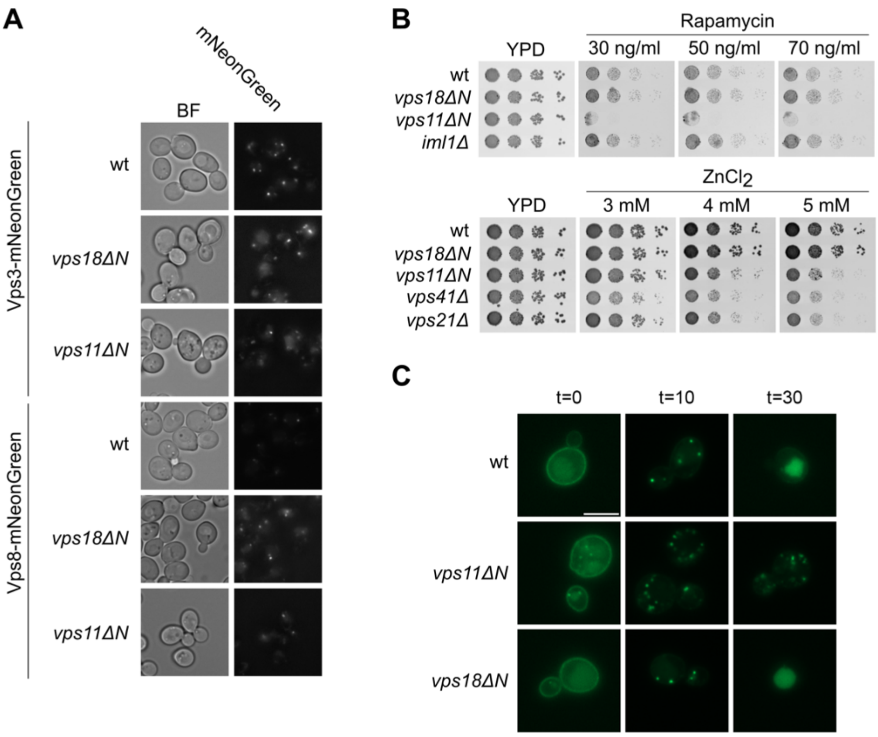


**Figure S8**. **Functional analysis of CORVET mutant strains. A**, Localization of Vps3 and Vps8 in Vps11 and *vps18ΔN* mutants. Vps3 or Vps8 were genomically tagged with mNeonGreen. Cells were grown to log phase and visualized by by fluorescence microscopy. **B**, Growth test assays. Cells were grown to the same $OD_{600}$ and spotted in serial dilutions on YPD without or with Rapamycin or $ZnCl_2$. Plates were grown at 30°C for 1-2 days before imaging. **C,** Mup1 uptake assay. Mup1 was C-terminally tagged with msGFP2 in wild-type, *vps11ΔN* and *vps18ΔN* cells. Cells were grown in synthetic medium lacking methionine and analyzed by fluorescence microscopy (t=0), before shifting to methionine containing medium for 10 min (t=10) and 30 min (t=30). Scale bar: 5 µm. Quantification is shown in Figure 5.

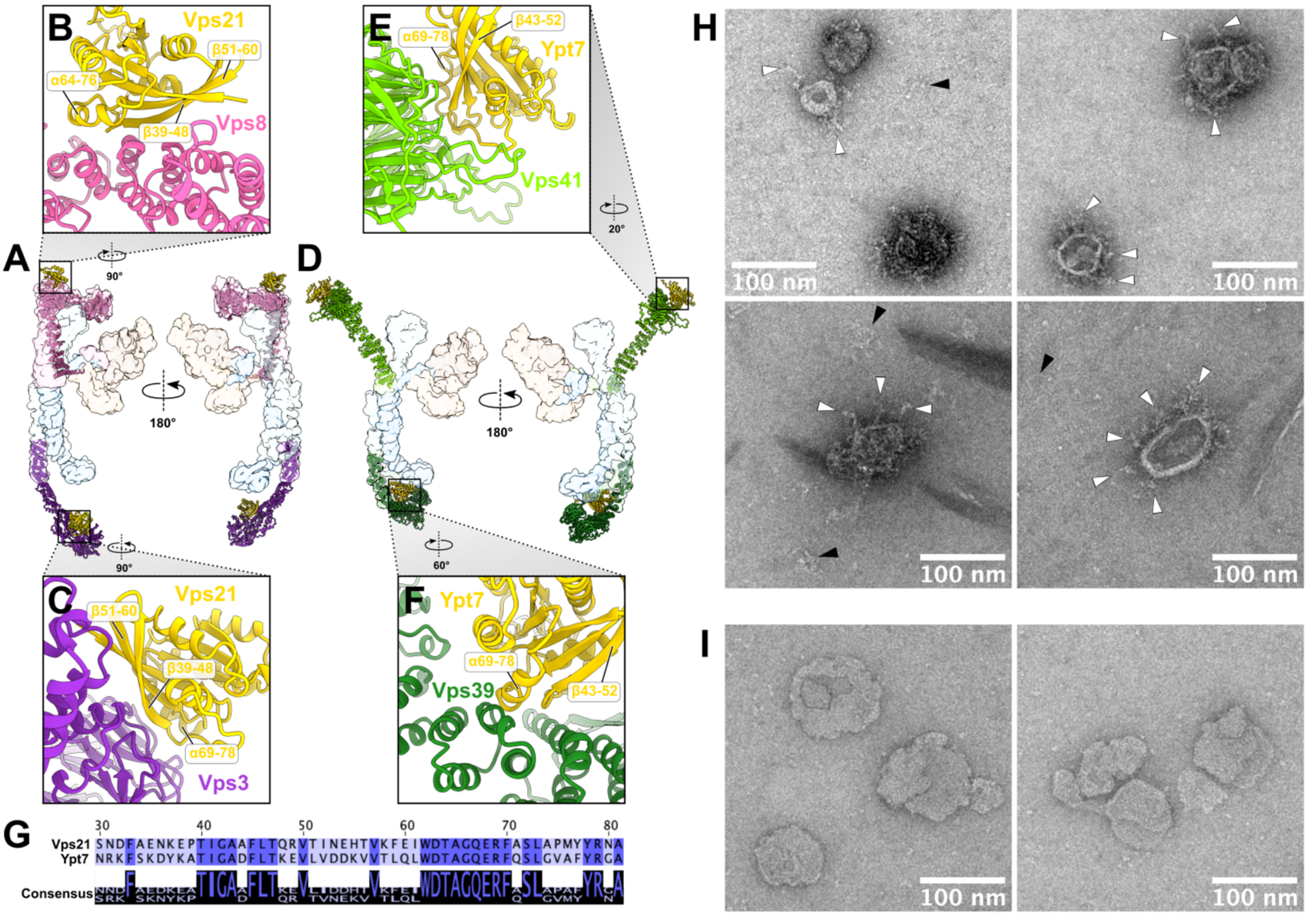


**Figure S9**. **Analysis of CORVET interactions with Rab GTPases and membranes. A,** Alphafold models Vps8 (pink) and Vps3 (violet) complexed with the Rab GTPase Vps21 (yellow) fitted into the structure of CORVET (this study, semi-transparent envelope) viewed from two sides. **B, C**, Close-up views at the predicted Vps8-Vps21 (**B**) and Vps3-Vps21 (**C**) interfaces. Secondary structural elements of Vps21 involved in the interface are indicated. **D**, Alphafold models Vps41 (light green) and Vps39 (dark green) complexed with the Rab GTPase Ypt7 (yellow) fitted into the structure of HOPS (PDB:7ZU0, semi-transparent envelope) viewed from two sides. **E, F**, Close-up views at the predicted Vps41-Ypt7 (**E**) and Vps39-Ypt7 (**F**) interfaces. Secondary structural elements of Vps21 involved in the interface are indicated. **G**, Sequence alignment of Vps21 and Ypt7 showing conservation of protein fragments predicted to be involved in the interaction (see **B, C, E, F**). **H,I,** Negative stain EM analysis of Rab5(Vps21)-loaded liposomes containing 10 mol % PI3P incubated with CORVET (**H**) or without (**I**).

### Supplementary Tables

**Table S1. Yeast strains used in the study.**

| Strain | Genotype | Source |
|---|---|---|
| SEY6210 | MAT alpha *leu2-3,112 ura3-52 his3-Δ200 trp-Δ901 lys2-801 suc2-Δ9 GAL* | Reggiori Lab |
| CUY2489 | MAT alpha *his3-Δ200 leu2-Δ0 lys2-Δ0 met15-Δ0 trp1-Δ63 ura3-Δ0 VPS11::HIS3-GAL1pr VPS16::NatNT2-GAL1pr VPS18::KanMX-GAL1pr* | Ostrowicz et al., 2010 |
| CUY13053 | MATa *his3-Δ200 met15-Δ0 trp1-Δ63 ura3-Δ0 VPS8::TRP1-GAL1pr VPS3::HIS3-GAL1pr VPS33::KanMX-GAL1pr VPS8::HphNT1-FLAG* | This study |
| CUY13050 | MATa *his3-Δ200 met15-Δ0 trp1-Δ63 ura3-Δ0 VPS41::TRP1-GAL1pr VPS39::KanMX-GAL1pr VPS33::HIS3-GAL1pr VPS41::HphNT1-FLAG* | Shvarev et al., 2022 |
| CUY13080 | CUY2489xCUY13050 | Shvarev et al., 2022 |
| CUY13081 | CUY2489xCUY13053 | This study |
| CUY14313 | MAT alpha *his3-Δ200 leu2-Δ0 lys2-Δ0 met15-Δ0 trp1-Δ63 ura3-Δ0 VPS11Δaa1-349::URA3-GAL1Pr VPS16::NatNT2-GAL1pr VPS18::KanMX-GAL1pr x*<br>MATa *his3-Δ200 leu2-Δ0 met15-Δ0 trp1-Δ63 ura3-Δ0 VPS8::TRP1-GAL1pr VPS3::HIS3-GAL1pr VPS8::HphNT1-FLAG VPS33::KanMX-GAL1pr* | This study |
| CUY14318 | MATa *his3-Δ200 met15-Δ0 trp1-Δ63 ura3-Δ0 VPS8::TRP1-GAL1pr VPS33::URA3-GAL1pr Vps3::HIS3-GAL1pr* | This study |
| CUY13750 | MAT alpha *his3-Δ200 leu2-Δ0 lys2-Δ0 met15-Δ0 trp1-Δ63 ura3-Δ0 VPS11::HIS3-GAL1pr VPS16::kanMX-GALpr1 VPS18Δaa1-335::TRP1-GAL1pr VPS18::FLAG-HphNT1* | Shvarev et al., 2022 |
| CUY14325 | CUY14318xCUY13750 | This study |
| CUY2742 | Mat alpha *vps11-1 leu2-3,112 ura3-52 his3-Δ200 trp1-Δ901 lys2-801 suc2-Δ9 VPS11-1::HIS3-HA* | Robinson et al., 1991 |
| CUY2743 | Mat alpha *vps11-3 leu2-3,112 ura3-52 his3-Δ200 trp1-Δ901 lys2-801 suc2-Δ9* | Robinson et al., 1991 |
| CUY2744 | Mat alpha *vps18-1 leu2-3,112 his3-Δ200 trp1-Δ901 lys2-801 suc2-Δ9* | Robinson et al., 1991 |
| CUY13541 | MAT alpha *leu2-3,112 ura3-52 his3-Δ200 trp-Δ901 lys2-801 suc2-Δ9 VPS8::ALFA-HphNT1* | This study |

| | | |
|---|---|---|
| CUY13599 | Mat alpha *leu2-3,112 ura3-52 his3-Δ200 trp1-Δ901 lys2-801 suc2-Δ9 VPS11-1::HIS3-HA VPS8::ALFA-HphNT1* | This study |
| CUY13558 | Mat alpha *leu2-3,112 ura3-52 his3-Δ200 trp1-Δ901 lys2-801 suc2-Δ9 VPS8::ALFA-HphNT1* | This study |
| CUY13560 | Mat alpha *leu2-3,112 his3-Δ200 trp1-Δ901 lys2-801 suc2-Δ9 VPS8::ALFA-HphNT1* | This study |

Ostrowicz, C. W. *et al*. Defined Subunit Arrangement and Rab Interactions Are Required for Functionality of the HOPS Tethering Complex. *Traffic* 11, 1334–1346 (2010).

Robinson, J.S. *et al*. A putative zinc finger protein, Saccharomyces cerevisiae Vps18p, affects late Golgi functions required for vacuolar protein sorting and efficient alpha-factor prohormone maturation. *Mol Cell Biol* 11(12), 5813-5824 (1991).

Shvarev, D. *et al*. Structure of the HOPS tethering complex, a lysosomal membrane fusion machinery. *eLife* vol. 11 e80901. 13 Sep. 2022, doi:10.7554/eLife.8090

**Table S2. Oligonucleotides used in the study.**

| Primer | Sequence | used for |
|---|---|---|
| S1 Vps3 | cagtcaaggagactacctttttttggttgcaaccataat attatagaaccgaattcgagctcgtttaaac | N-terminal tagging with Gal1 promotor |
| S4 Vps3 | ccttcattttcctttacttcttttcctttgtcattattcgttttctttt ttaccattttgagatccgggtttt | N-terminal tagging with Gal1 promotor |
| S1 Vps8 | ggcttaataagtgtaaaatatatatctgccgagaccatt actcattacacctagagaattcgagctcgtttaaac | N-terminal tagging with Gal1 promotor |
| S4 Vps8 | gtcgtatcgatgctagatctgctgtcgtggtcaaggcca ttttgctccattttgagatccgggtttt | N-terminal tagging with Gal1 promotor |
| S1 Vps11 | agttaaaccttccaaaaatcatagcgtttcatctatagg cacagcaaatcagtcgtacgctgcaggtcgag | N-terminal tagging with Gal1 promotor |
| S4 Vps11 | cttatgggaatattctcgaaaagctggaattgcctcca ggagctcagggacatcgatgaattctctgtct | N-terminal tagging with Gal1 promotor |
| S1 Vps16 | gaatagtacagacggataggcctcccttttgtttcgcac taataaaaatgcgtacgctgcaggtcgac | N-terminal tagging with Gal1 promotor |
| S4 Vps16 | ctataaaatacgtctttcaatctttcccagtcgaagctag ggtttttcatcgatgaattctctgtcg | N-terminal tagging with Gal1 promotor |
| S1 Vps18 | aaaaactataaggtaccaagaagtaaaaagagaa atatagggatataatgcgtacgctgcaggtcgac | N-terminal tagging with Gal1 promotor |
| S4 Vps18 | gtattccctgtgaggaattgtaactgaacttcctctatac gtgtttttatcatcgatgaattctctgtcg | N-terminal tagging with Gal1 promotor |
| S1 Vps33 | gaaaaagctgatattgcccatctcaaactttatcaaatc atttcacgatgcgtacgctgcaggtcgac | N-terminal tagging with Gal1 promotor |
| S4 Vps33 | gtccatcggcatttgttaatgaaaatttcttagtattccaa aatctattcatcgatgaattctctgtcg | N-terminal tagging with Gal1 promotor |
| S2 Vps8 | tataaattttacttttatgtaaccaaagttgtattaaatattt agaaatgatcgatgaattcgagctcg | C-terminal tagging with FLAG or ALFA tag |
| S3 Vps8 | atgaatattcttgtttaatttgccagacggaatctaaccc aaaaatagtacgtacgctgcaggtcgac | C-terminal tagging with FLAG or ALFA tag |
| S2 Vps18 | ccctcttttaatttcagtgtttcagcctgactaaaaaaga atactaatcgatgaattcgagctcg | C-terminal tagging with FLAG tag |
| S3 Vps18 | cagccaatatctattgatgaaacagaattagccaaat ggaatgaacgtacgctgcaggtcgac | C-terminal tagging with FLAG tag |
| S4 Vps18 aa336-end | aatttgttatcccgaactaataaattccataccgaatta ggctcattttccattttgagatccgggtttt | Truncation of Vps18 N-Terminus |
| S4 Vps11 aa349-end | tccttttggatgatgatgttaatttgattttctaaagatttcat atcatcgatgaattctctgtcg | Truncation of Vps11 N-Terminus |
| S2 Mup1 | gttcatacgtgattataagaatcgagatgagatggtaa gtacctttttggttaatcgatgaattcgagctcg | C-terminal tagging of Mup1 |
| S3 Mup1 | cgttattgaaacgaatataatcgaacattacaaaagtg aacaagaaaaatcgctgcgtacgctgcaggtcgac | C-terminal tagging of Mup1 |

**Table S3. Cryo-EM data collection, refinement and validation statistics.**

| | CORVET composite map (EMDB-xxxx) (PDB xxxx) | CORVET consensus map (EMDB-xxxx) | CORVET Vps8-Vps11 local (EMDB-xxxx) | CORVET Vps16-Vps33 local (EMDB-xxxx) | CORVET Vps8 β-propeller local (EMDB-xxxx) | CORVET Vps18-Vps11 local (EMDB-xxxx) | CORVET Vps18 β-propeller local (EMDB-xxxx) |
|---|---|---|---|---|---|---|---|
| **Data collection and processing** | | | | | | | |
| Magnification | | | | 130,000 | | | |
| Voltage (kV) | | | | 200 | | | |
| Electron exposure (e–/Å$^2$) | | | | 50 | | | |
| Defocus range (μm) | | | | -0.8 to -2.8 | | | |
| Pixel size (Å) | | | | 0.924 | | | |
| Symmetry imposed | | | | C1 | | | |
| Initial particle images (no.) | | | | 1,445,679 (after duplicates removal) | | | |
| Final particle images (no.) | | 219,391 | 218,807 | 211,131 | 219,391 | 218,807 | 219,391 |
| Map resolution (Å) | 3.8-4.6 | 4.5 | 4.2 | 3.8 | 4.4 | 4.3 | 4.6 |
| FSC threshold | 0.143 | 0.143 | 0.143 | 0.143 | 0.143 | 0.143 | 0.143 |
| **Refinement** | | | | | | | |
| Initial model used | AlphaFold/ PDB:7ZU0 | - | - | - | - | - | - |
| Model resolution range (Å) | 3.8-4.6 | - | - | - | - | - | - |
| FSC threshold | 0.143 | | | | | | |
| Map sharpening *B* factor (Å$^2$) | - | 133.6 | 95.9 | 95.6 | 89.2 | 118.1 | 138.1 |
| Model composition | | | | | | | |
| Non-hydrogen atoms | 30231 | - | - | - | - | - | - |
| Protein residues | 4468 | | | | | | |
| R.m.s. deviations | | | | | | | |
| Bond lengths (Å) | 0.002 | - | - | - | - | - | - |
| Bond angles (°) | 0.554 | | | | | | |
| Validation | | | | | | | |
| MolProbity score | 2.03 | - | - | - | - | - | - |
| Clashscore | 8.74 | | | | | | |
| Poor rotamers (%) | 0 | | | | | | |
| Ramachandran plot | | | | | | | |
| Favored (%) | 89.68 | - | - | - | - | - | - |
| Outliers (%) | 0.07 | | | | | | |

#### Supplementary Movies

**Movie S1. Overall architecture of CORVET tethering complex.** Movie representing transition between molecular surface and ribbon representation of the structure. Coloring as in Figure 1.

**Movies S2-S4. 3D variability of CORVET.** Movies show eigenvectors of variability in the dataset (variability components 000-002, also see Figure S5).
